# Orphan nuclear receptor *NR2E3* and its small-molecule agonist induce cancer cell apoptosis through regulating p53, IFNα and MYC pathways

**DOI:** 10.1101/2023.12.26.573366

**Authors:** Yidan Wang, Todd Kroll, Linhui Hao, Ansul Sharma, Vivian Zhou, Luke Moat, John Mayer, Sanjay S. Shukla, Scott Hebbring, Song Guo, Marissa Iden, Adam Bissonnette, Gene Ananiev, Deepak Parashar, Janet S. Rader, Siegfried Janz, Zhi Wen

## Abstract

Orphan nuclear receptor NR2E3 activates p53 and induces cancer cell apoptosis. Further studies on p53-dependent and -independent functions of wild-type and mutated *NR2E3* are needed. Herein, we showed that NR2E3 enhanced p53-DNA interactions in diverse cancer cells and up-regulated p53 and IFNα pathways while down-regulating MYC pathway in cervical cancer cells. Studies of “All *of* Us” and TCGA databases showed *NR2E3* nonsynonymous mutations’ associating with four cancers. We stratified *NR2E3* SNVs for their cancer implications with the p53 reporter. A cancer-associated *NR2E3^R97H^*mutation not only lost the wild-type’s tumor-suppressing functions but also prohibited the wild-type from enhancing p53 acetylation. These observations implicated the potential for pharmaceutically activating NR2E3 to suppress cancer. Indeed, NR2E3’s small-molecule agonist 11a repressed 2-D and 3-D cultures of primary cells and cell lines of cervical cancer, in which screening FDA-approved anti-cancer drugs identified HDAC-1/2 inhibitor Romidepsin operating synergistically with 11a. The underlying molecular mechanisms included 11a’s down-regulating the transcription of Multidrug Resistance Protein *ABCB1* that Romidepsin up-regulated. Transcriptomics studies revealed three synergy modes: (1) “sum-up” mode that the p53 pathway activated individually by 11a and Romidepsin got stronger by the combo; (2) “antagonism” mode that Romidepsin counteracted the activation of the Kras pathway by 11a; and (3) “de novo” mode that the combo instead of each individual drug repressed the MYC pathway. Conclusively, our experiments provide new data supporting tumor-suppressor like functions for wild-type *NR2E3*, reveal roles of mutated *NR2E3* in cancer, and address values of NR2E3’s agonist 11a in cancer therapy alone and combined.

## Introduction

Orphan Nuclear Receptor subfamily 2 group E member 3 (*NR2E3*) was first cloned from human retinoblastoma Y79 cells (1) and a mouse λZAP eye cDNA library (2) in 1999. The *NR2E3* gene locates between q22.33-->q23 on chromosome-15 in human and on chromosome-9 in mouse (3). There are two protein isoforms of NR2E3 in human and one isoform in mouse. Human full-length NR2E3 consists of 410 amino acids forming a DNA-binding domain (DBD Cys47-Val130) and a Ligand-binding domain (LBD His221-Asn410) (4). The crystal structure of the LBD displays a dimeric auto-repressed conformation (5). In contrast, the short isoform lacks the C-terminal 43 amino acids. NR2E3 is not only a transcription factor that binds to a consensus DNA duplex site (AAAGTCAAAAGTCA) (1,6) but also is a transcriptional cofactor (7). NR2E3 is an orphan nuclear receptors whose natural ligands remain unknown. However, compounds with a 2-phenylbenzimidazole core, such as 11a, have been reported to be potent agonists of NR2E3 transactivity (8).

*NR2E3* mutations cause several inherited retinal degenerations such as Enhanced S-cone syndrome (9). The deletion of a 380-nt fragment in the mouse *Nr2e3* coding region damages the DBD domain and results in similar retinal degenerations in the *rd7* mouse model (10). These diseases are characterized by loss of rod photoreceptors and gain of cone photoreceptors (11). In this setting, *NR2E3* mutations may lose transcriptional activities that support the differentiation of rod photoreceptors. For example, *NR2E3^R97H^*mutant proteins fail to bind the NR2E3 consensus sequence (12). However, there is a unique later-stage dysplastic appearance in *NR2E3*-mutated retinal degenerations in human that raises the possibility that NR2E3 is involved in cell proliferation (13). The loss of *Nr2e3* causes retinal dysplasia (10) and increase in a p53-repressed gene *Survivin* (14) in *rd7* mice, suggesting that *NR2E3* may also regulate cell survival.

We previously identified NR2E3 as a host factor activating p53 in Human Papilloma Virus (HPV)-positive cervical cancer cells, following a high-throughput screen of 16,000 human and mouse cDNAs (15). p53 is a key tumor suppressor that induces cell apoptosis among other functions (16-18). Loss of functional p53 has been detected in nearly all cancers, mainly due to mutations and dysregulated post-translational modifications (*e.g.,* acetylation) (19,20). For instance, a chief role of the high-risk HPV oncoprotein E6 in cervical cancer carcinogenesis is stimulation of p53 ubiquitination and proteasomal degradation (21). We have shown that NR2E3 serves as a transcriptional cofactor of p53 by stimulating p300/PCAF-mediated acetylation of p53 and cell apoptosis in HPV^+^ cervical cancer cells (15). We further discovered p53-enhanced cell-killing effects by NR2E3’s agonist 11a in screening the NCI-60 cancer cell panel (22). Other colleagues have shown positive associations between NR2E3 and recurrence-free survival in ER^+^ breast cancer (23) and overall survival in liver cancer (24), and have identified p53 activation by NR2E3 through a long non-coding RNA (25). Taken together, these findings suggest a tumor suppressive function for *NR2E3* that is incompletely understood. Herein, we further determined molecular roles of wild-type and mutated *NR2E3* and identify *NR2E3*-targeted combinatorial therapy that may be applicable for cancer.

## Methods and Materials

Please see the supplemental files.

## Results

### Full-length NR2E3 regulates p53, IFNα and other growth-related pathways in cancer cells

Universal expression of NR2E3 has been detected in urogenital and respiratory systems in addition to retinal photoreceptor cells in human and mouse (supplementary Fig. S1). To further our understanding of NR2E3’s roles in cancer, we expressed HA-tagged NR2E3 proteins of full-length isoform (FL), DNA binding domain (DBD), ligand binding domain (LBD) and short isoform (short) in HeLa cervical cancer cells. FL and LBD localized inside the nucleus, whereas short and DBD localized outside the nucleus (Fig. 1B), indicating that a nuclear localization signal resides in the 43 amino acids at the C-terminus of NR2E3.

**Figure 1:**
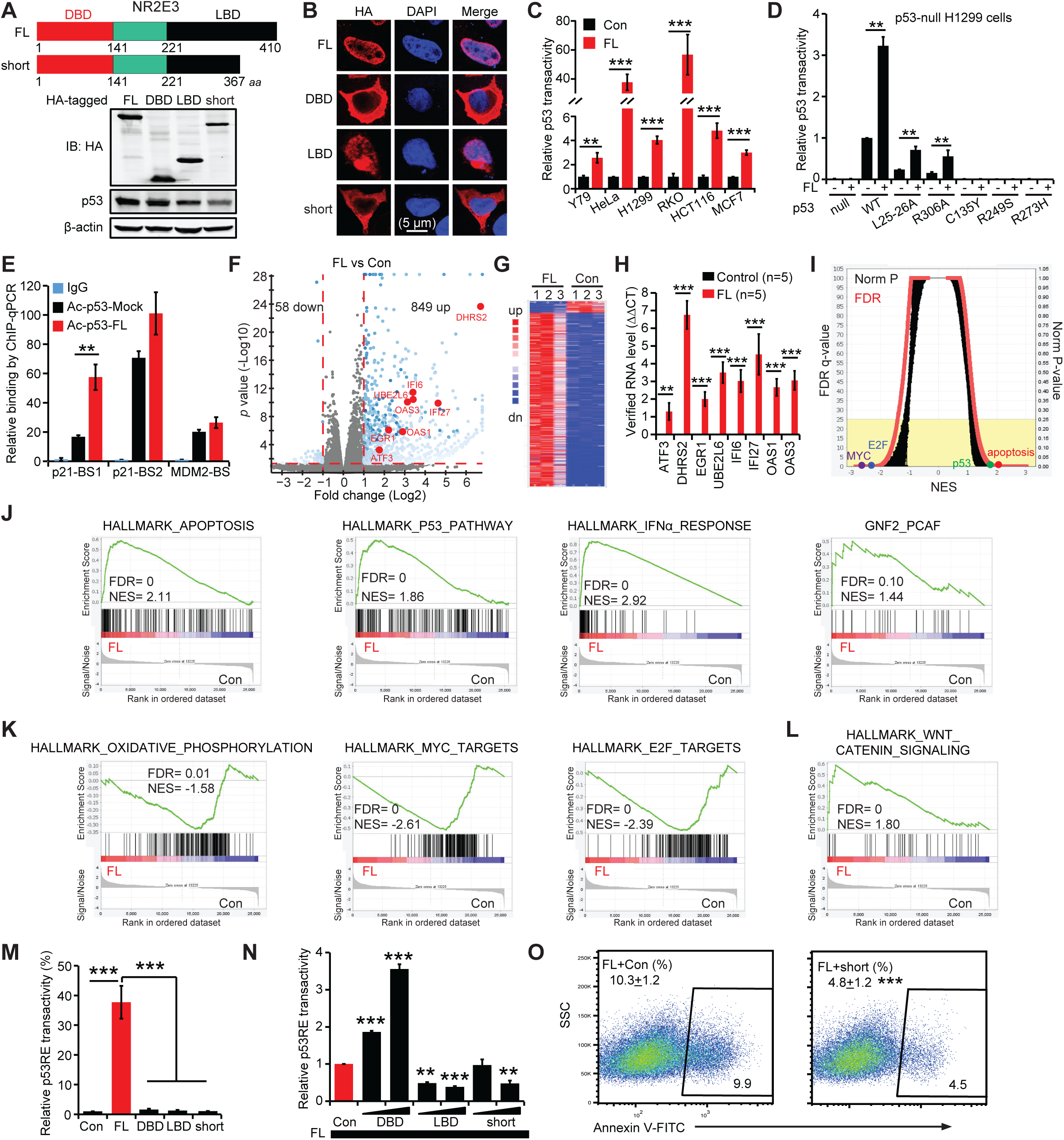
NR2E3’s short isoform abrogates its full-length isoform from suppressing cancer cells. (A) The short failed to increase p53 protein in HeLa cells. *Up*: illustration of human NR2E3 proteins. DBD: DNA binding domain; LBD: ligand binding domain; FL: full-length isoform; and short: short-length isoform. *Below*: immunoblotting analysis of HA-tagged NR2E3 constructs and p53. (B) The short lost nuclear localization signal. Immunofluorescence with anti-HA antibody was conducted in HeLa cells. (C) The FL activated a p53-responsive luminescence reporter in 6 cancer cell lines. These cells endogenously express wild-type p53, except H1299 cells with enforced p53 expression. The activity in mock control was normalized to 1. (D) The FL selectively rescued the wild-type transactivities of p53 mutations partially reserving this transactivity in H1299 cells. The activity in the p53^WT^+empty vector control was normalized to 1. (E) The FL enhanced acetylated p53’s binding to p21 promoter. Anti-acetylated p53 antibody was used in chromatin immunoprecipitation (ChIP) followed by qPCR analysis. *BS:* p53-binding site. (F-L) RNA sequencing analysis of HeLa cells enforcedly expressing the FL or mock control. (F) Volcano plot of all transcripts. *Red:* growth inhibitory genes. (G) Heat map of differentially expressed genes. (H) Up-regulations of growth inhibitory genes were verified by qRT-PCR. (I) Overview of GSEA results. (J) Gene sets of p53, apoptosis, INFα and PCAF were enriched in FL group. (K) Metabolic oxidative phosphorylation pathway and oncogenic MYC and E2F pathways were enriched in control group. (L) Oncogenic WNTβ pathway was enriched in FL group. (M) The short failed to activate the p53 reporter in HeLa cells. (N) The short inhibited the FL’s activation of the p53 reporter in HeLa cells. The activity in the FL+empty vector control was normalized to 1. (O) The short inhibited the FL-induced HeLa cell apoptosis in Annexin V-binding assay. *p* value was calculated by Student’s t-test with two tails. *: *p*<0.05; **: *p*<0.01; ***: *p*<0.001.

Stimulation of p53 transactivity (p53-mediated transactivation) by FL was confirmed in HeLa cells in addition to human retinoblastoma-Y79, lung cancer-H1299, colon cancer-RKO and HCT116, and breast cancer-MCF7 cells all of which express wild-type p53 (Fig. 1C), showing a conserved cellular machinery for NR2E3’s activating p53 in > 5 cancer cell types. To test whether mutations in p53 affect this NR2E3 regulation, a p53 mutation L25-26A that damages its Transactivation Domain (26), a dominant-negative mutation C135Y, a hot-spot mutation R249S, a gain-of-function mutation R273H, and R306A that disrupts nuclear localization of p53 (27), were co-expressed with FL in the p53 reporter assay. p53^C135Y^, p53^R249S^ and p53^R273H^ completely lost the ability to stimulate p53 transactivity in *p53^-/-^* H1299 cells and *p53^-/-^* HCT116 cells, whereas p53^L25-26A^ and p53^R306A^ reduced p53 transactivity by 70-80% (Fig. 1D & supplementary Fig. S2). FL did enhance the wild-type transactivities of p53^L25-26A^ and p53^R306A^ instead of p53^C135Y^, p53^R249S^ and p53^R273H^ (Fig. 1D & supplementary Fig. S2), implying that NR2E3 can activate these p53^MUT^ with residual wild-type transactivity. Interestingly, related subfamily members *NR2E1*, *NR2F1* and *NR2F2* also stimulated p53 transactivity in HeLa and HCT116 cells (supplementary Fig. S3), showing a common role for NR2 members in regulation of p53.

NR2E3 enhances acetylation of p53 and this selectively up-regulates some gene targets (*e.g.,* p21) compared to others (*e.g.,* MDM2) (19). As shown in Figure 1E, FL enhanced acetylated p53’s binding to its DNA consensus in the *p21* gene but not in the *MDM2* gene in HeLa cells. To further explore NR2E3 functions in gene expression, we conducted an RNA sequencing study in HeLa cells (Fig. 1F-L). FL up-regulated 849 genes and down-regulated 58 genes (fold change > 2; *p* < 0.05) (Fig. 1F-G). Up-regulation included 8 genes that induce cell apoptosis or inhibit cell growth, and they were verified by qRT-PCR (Fig. 1H). For example, UBE2L6 encodes a ubiquitin-conjugating enzyme UBCH8 which redirects E6AP to the ubiquitination of HPV E6-independent substrates (28). UBE2L6 is also a p53-responsive gene under DNA damage conditions (29). IFI6, IFI27, OAS1 and OAS3 are genes in the interferon-alpha (IFNα) response pathway which mediates anti-tumor and anti-viral immune responses (30). Gene set enrichment analysis (GSEA) confirmed up-regulated apoptosis, p53, IFNα, and p53-acetyltransferase PCAF pathways by FL (Fig. 1I-J). In contrast, FL repressed Oxidative Phosphorylation-related ATP production and oncogenic MYC and E2F pathways (Fig. 1K). However, these cells tried to alleviate cell apoptosis by enhancing WNT signaling (31) (Fig. 1L), implicating a demand of combinatorial therapy to suppress survival signals developed by activated NR2E3. Together, these data show that full-length NR2E3 regulates p53 and other cancer pathways such as IFNα and MYC.

The short isoform of NR2E3 arises naturally from alternative mRNA splicing. In contrast to FL, short and DBD were unable to stimulate p53 transactivity (Fig. 1M), likely because of their cytosolic localizations. LBD neither stimulated p53 transactivity, possibly because of its deficit in binding DNA (Fig. 1M). Interestingly, both LBD and short prevented FL from stimulating p53 transactivity when co-expressed with FL, whereas DBD increased FL-stimulated p53 transactivity under the same conditions (Fig 1N). When co-expressed with FL, short inhibited cell apoptosis over 2-fold (Fig. 1O). These data suggest that a balance of natural NR2E3 isoforms likely determines overall NR2E3 effects in cancer.

### *NR2E3* is associated with cancer incidence and prognosis

Over 500 single nucleotide variants (SNV) have been identified in *NR2E3* in ClinVar and many have been associated with genetic diseases. These *NR2E3* SNVs are all germline ones which led to 272 amino acids changes (supplementary Fig. S4A). Most of the SNVs have been reported in clinical specimens without follow-up studies except the predictions to encompass “benign”, “pathogenic”, and “uncertain significance” mutations (supplementary Fig. S4B).

TCGA database carries the mutations data in cancer patient cohorts, which can be used to evaluate the association between mutation frequency and cancer incidence when a reference database of regular population is available. However, the frequencies of *NR2E3* SNVs are extremely low in regular population. Taking advantage of being a part of the NIH “All *of* Us” research program, we surveyed the “All *of* Us” database that collected comprehensive information of > 182,000 cases representing the diversities of race, age, sex, health status, income, geography, lifestyle, *etc.* as the reference of TCGA cancer cases. Nonsynonymous SNVs of *NR2E3* were counted in both cancer cases and reference population. 11/522 (*NR2E3* mutation *vs* WT cases) in colon cancer patients, 5/954 in multiple myeloma patients, 19/733 in uterus cancer patients and 8/462 in melanoma patients were observed (Fig. 2A). There were significant associations between *NR2E3* SNVs and these 4 cancer types, compared to the reference population (*p* < 0.05 and high OR with 95% CI after considering race, sex and age), suggesting that *NR2E3* nonsynonymous SNVs are a risk factor for these cancer types (Fig. 2A).

**Figure 2:**
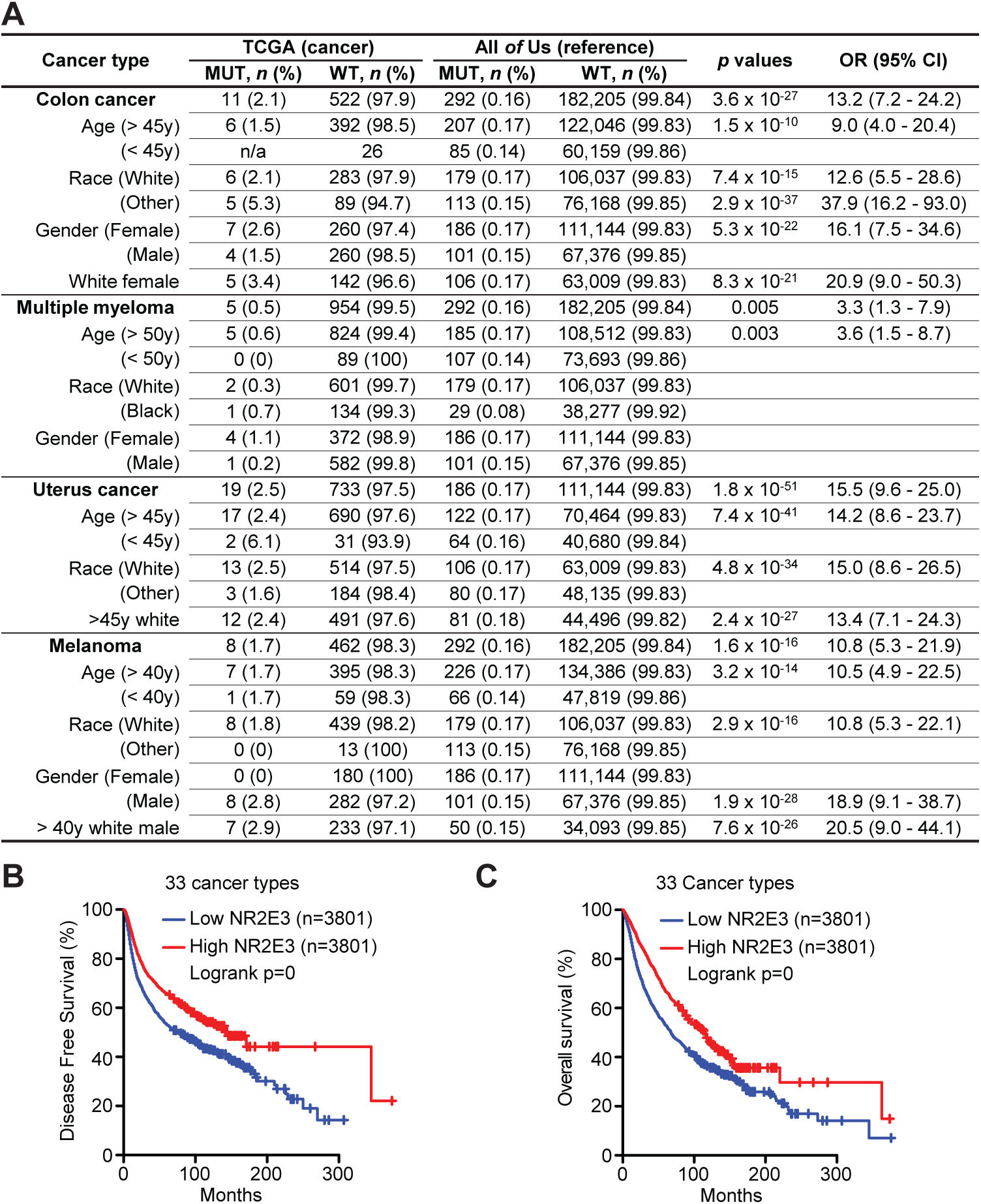
NR2E3 is closely associated with cancer. (A) NR2E3 mutations were enriched in colon cancer, multiple myeloma, uterus cancer and melanoma. Nonsynonymous SNVs of NR2E3 were counted in TCGA (incorporating MMRF) cancer database and “All *of* Us” reference database. Age, sex and race were considered. The age group cutoffs were set (45-47). Chi-square test (two-tail, 1 df) and Woolf logit test (n > 99k) or Baptista-Pike (n < 99k) test were used to calculate the *p* values and OR values with 95% confidence interval only in the comparisons with minimal case number of 5. (B-C) High expression levels of NR2E3 were correlated to superior Disease-free cancer survival (B) and overall survival (C) of 33 cancer types. GEPIA tool was used to analyze the data in TCGA database. The group cutoff: High is > 60%; and low is < 40%. NR2E3 level was normalized to β-actin. *p* values were calculated by Logrank test.

Interestingly, the statistic tool GEPIA (32) also linked *NR2E3* mRNA levels to overall and disease-free survival of cancer patients in the TCGA database. High *NR2E3* RNA levels were associated with extended disease-free (Fig. 2B) and overall (Fig. 2C) survivals for 33 cancer types combined, further supporting the hypothesis that *NR2E3* has a tumor suppressor like function. Associations between *NR2E3* expression levels and prognosis in breast and liver cancers have also been reported (23-25), reaffirming that high expression of *NR2E3* predicts superior cancer prognosis.

### Stratify *NR2E3* SNVs with the p53 reporter

Stratifying > 500 *NR2E3* SNVs helps to understand their roles in carcinogenesis. The p53 reporter assay could be a simple reliable tool for this task. We selected 6 of the 25 disease-associated SNVs (supplementary Fig. S5) for structure-function studies in cancer cells as they were previously studied in human and monkey kidney cells (12). ClinVar suggested R76W, G88V and R97H in *NR2E3* as pathogenic mutations, E121K and V302I as benign, and M407K has not yet been annotated. Interestingly, 2 of 512 uterus endometrial cancers in TCGA database also harbor the R97H mutation. Though higher than other tissues, *NR2E3* expression is still low in urogenital system. Therefore, we adopted enforced-expression assays to visualize the differences in activating p53 between *NR2E3* SNVs, instead of Crispr/Cas9 tool to knock in point mutations in cells to study them at a physiological state.

Compared to FL, R76W, G88V and R97H did not activate the p53 reporter in HeLa, HCT116 and H1299 cells (Fig. 3A, supplementary Fig. S6), supporting the idea that they are “pathogenic” for p53-related functions of wild-type *NR2E3*. In contrast, E121K, V302I and M407K retained the ability to active the p53 reporter, behaving “benign”. Expression of R76W, G88V and R97H did not increase p53 protein expression, whereas E121K, V302I and M407K increased p53 protein levels (Fig. 3B). Interestingly, R76W, G88V and R97H did not enhance cell apoptosis in contrast to FL, whereas E121K and V302I had some variable apoptosis effects (Fig. 3C). In summary, pathogenic *NR2E3* mutations, including the R97H mutation that was detected in endometrial cancer tissues, but not benign or uncertain ones lose the ability to enhance p53 transactivity and cell apoptosis. Our results indicate the p53 reporter assay as a simple reliable experimental tool to stratify *NR2E3* SNVs with implications in cancer.

**Figure 3:**
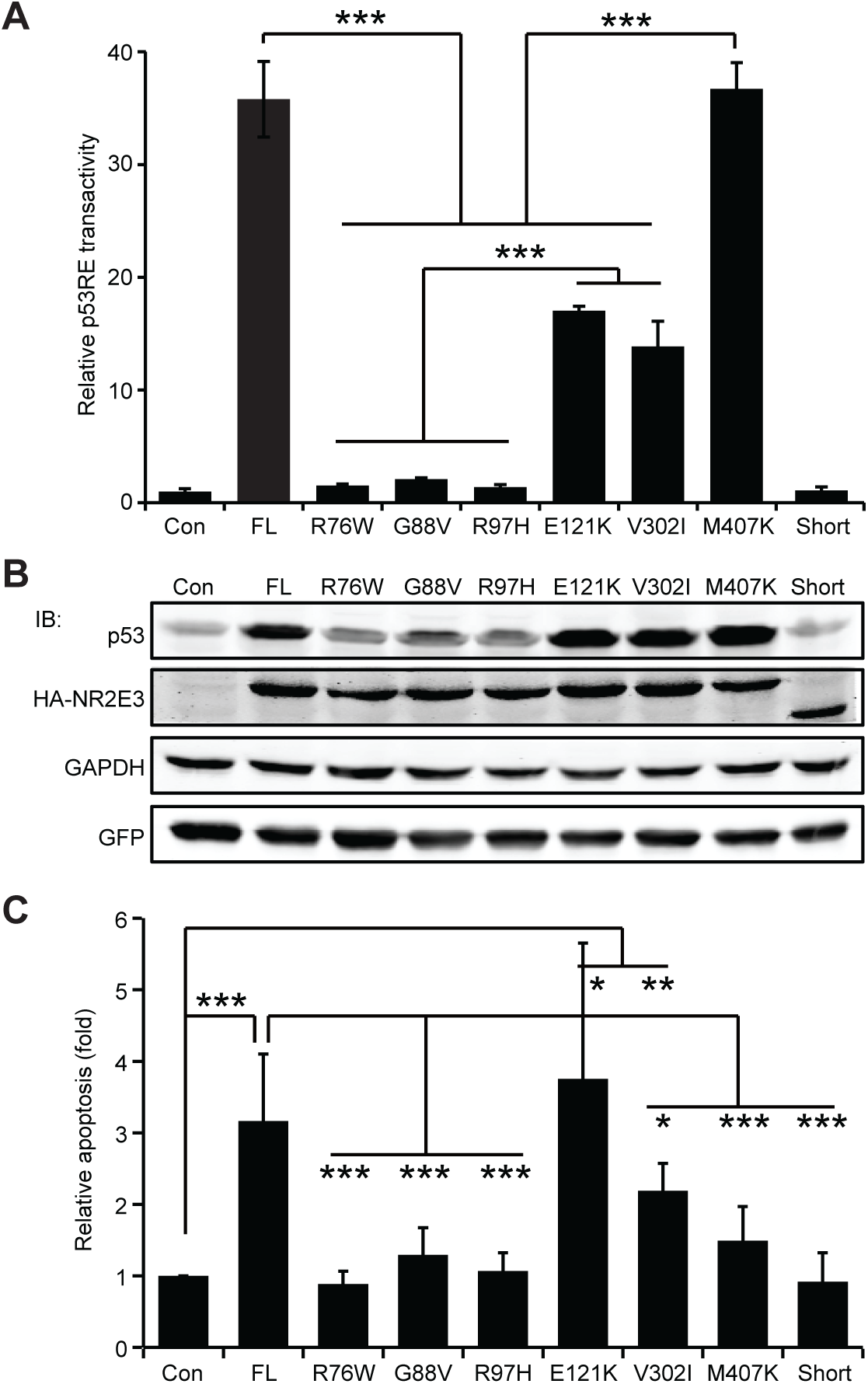
NR2E3 SNVs are stratified with their abilities to activate p53. (A) NR2E3 SNVs differentially stimulated the p53 reporter in HeLa cells. The activity in control was normalized to 1. (B) NR2E3 SNVs differentially regulated p53 protein levels in HeLa cells in immunoblotting assay. GFP: transfection reference. (C) NR2E3 SNVs differentially regulated HeLa cell apoptosis in Annexin V-binding assay. The apoptosis in the mock control was normalized to 1. *p* value was calculated by Student’s t-test with two tails. *: *p*<0.05; **: *p*<0.01; ***: *p*<0.001.

### R76W and R97H mutations lose the tumor-suppressing functions of wild-type *NR2E3*

We examined further the functional activities of the R76W and R97H mutations. Unlike FL, R76W and R97H were reported to localize to both the cytoplasm and the nucleus in HEK293 cells (12). We observed a similar subcellular localization of R76W and R97H in HeLa cells, but with less co-localization between p53 and R76W or R97H, compared to FL (Fig. 4A). Anti-p300 antibody co-immunoprecipitated FL, R76W and R97H to similar degrees, showing that these mutations do not disrupt their associations with p300 (Fig. 4B). Whereas FL increased the association between p300 and p53 (15), R76W and R97H had a reduced p300-p53 association (Fig. 4B). In addition, the extended half-life time of p53 in association with FL was lost in the R76W and R97H groups (Fig. 4C). In fact, a Gel-shift assay with ^32^P-labeled probe containing classic p53-binding consensus sequences showed that FL boosted p53-probe binding in a dose-dependent fashion, whereas R76W and R97H did not (Fig. 4D). Our data show that R76W and R97H are loss-of-function in their abilities to bind and activate p53.

**Figure 4:**
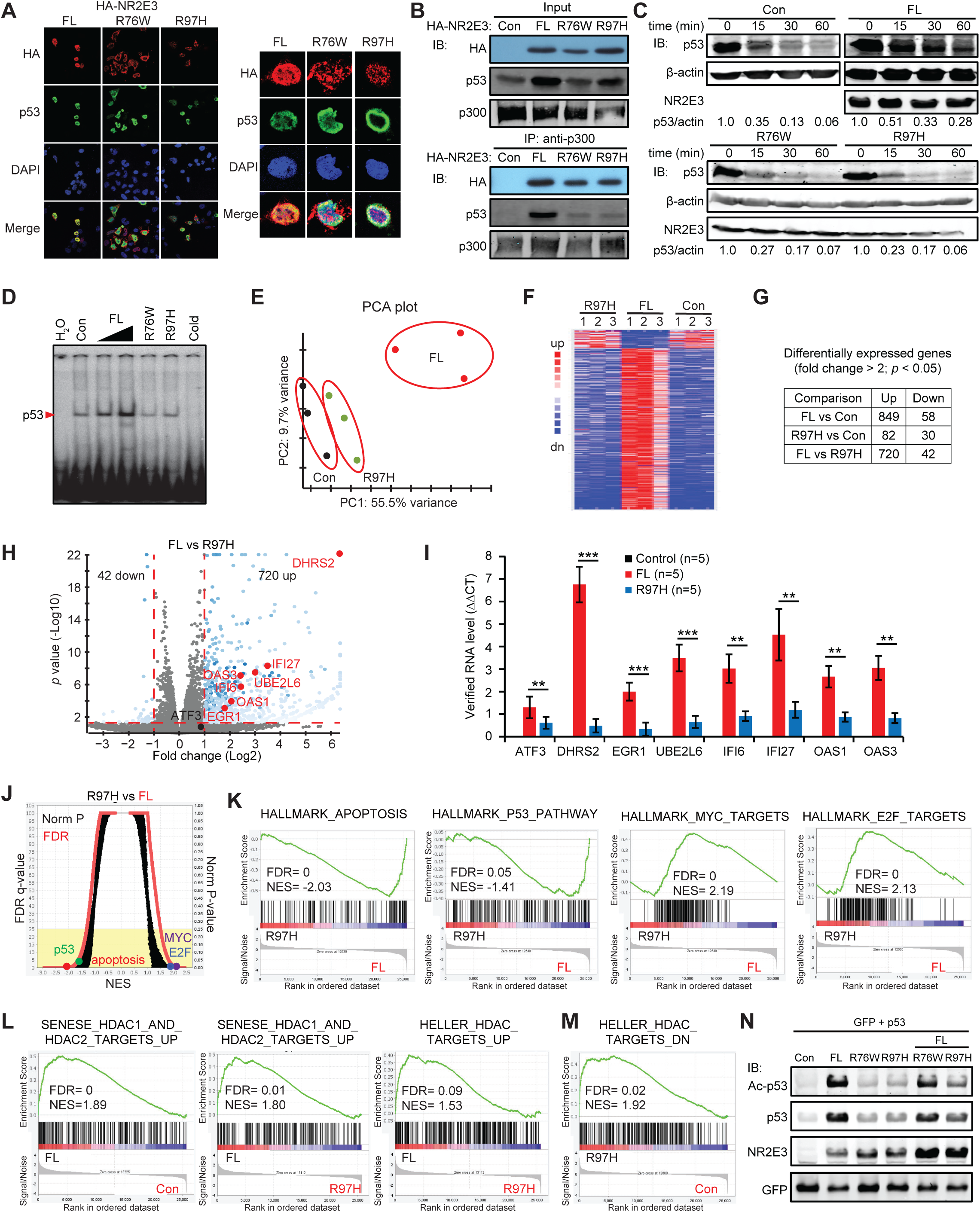
NR2E3 R76W and R97H are loss of function mutations. (A) Immunofluorescence analysis of co-localization of p53 and FL/R76W/R97H in HeLa cells. Zoom-in images for the details. (B) R76W and R97H failed to enhance p300’s binding to p53 in co-immunoprecipitation assay with an anti-p300 antibody followed by immunoblotting analysis. (C) R76W and R97H failed to extend half-life time of p53 protein in HeLa cells. The protein synthesis was ceased by Cycloheximide. (D) R76W and R97H failed to enhance HeLa cell nuclear extract’s binding to p53-DNA consensus in gel shift analysis. Cold (unlabeled) probe was loaded 50-fold more than the labeled probe. (E-M) RNA sequencing analysis of HeLa cells enforcedly expressing the FL, R97H or mock control. (E) Principal Component Analysis (PCA) plot of 9 samples. (F) Heat map of differentially expressed genes in FL, R97H and control groups. (G) The table summarized the number of differentially expressed genes in each comparison. (H) Volcano plot of the transcripts of FL and R97H groups. *Red:* growth inhibitory genes. (I) Up-regulations of growth inhibitory genes were verified using qRT-PCR. (J) Overview of GSEA results in R97H vs FL. (K) p53 and apoptosis pathways were enriched in FL group while MYC and E2F pathways were enriched in R97H group. (L) Acetylation-related gene sets were enriched in FL group. (M) Deacetylation-related pathway was enriched in R97H group. (N) R97H abolished the FL from acetylating p53.

We next explored the roles of the cancer-associated pathogenic R97H mutation in transcriptomics studies. The RNA expression pattern induced by R97H was similar to mock controls and dramatically different from FL (Fig. 4E-H). For example, 8 genes in p53 and IFNα pathways that were up-regulated by FL were largely unaffected by R97H (Fig. 4I). Compared to FL, R97H also failed to enhance the apoptosis and p53 pathways or inhibit MYC and E2F pathways (Fig. 4J-K), demonstrating that the R97H mutation deregulates p53, apoptosis and other important cell growth and cancer-related pathways

More specifically, whereas FL enriched gene sets that were up-regulated by RNAi- or Trichostatin A-induced inhibition of histone deacetylases (HDAC) (Fig. 4L), R97H increased genes that were down-regulated by Trichostatin A (Fig. 4M), suggesting that R97H may have a role in HDAC function. This presumptive role of R97H in HDAC function was further supported by the observation that R97H as well as R76W failed to enhance p53 acetylation and that R97H inhibited FL-mediated p53 acetylation (Fig. 4N). Collectively, our data show that R76W and R97H mutations in NR2E3 are loss-of-function in their ability to regulate p53 and other cancer pathways. Our studies of wild-type and mutated *NR2E3* suggest that pharmaceutically activating NR2E3 may contribute to cancer therapy.

### NR2E3’s agonist-11a synergizes with Romidepsin to suppress cervical cancer cells and spheroids

Compound-11a with a 2-phenylbenzimidazole core has been identified as an agonist of NR2E3 transcriptional activity (8). We previously reported that cells expressing wild-type p53 are more sensitive to growth inhibition by 11a than cells lacking wild-type p53 signaling (22). For example, *p53^+/+^* HCT116 cells were 100-fold more sensitive to 11a than their *p53^-/-^* counterpart (supplementary Fig. S7). We conducted drug repurposing screens of a FDA-approved anti-cancer drug library (NCI AOD X version) in the presence and absence of 11a to identify drugs that cooperate with 11a in killing HeLa cervical cancer cells (Fig. 5A). Romidepsin, a potent HDAC-1/2 inhibitor (HDACi), boosted 11a’s killing effects while Pralatrexate, a folate analog, antagonized 11a (Fig. 5A). Extra HDACi and DNA intercalating agents also boosted the cell-killing effects of 11a (*data not shown*). Five of these agents, including Romidepsin, have been used or tested in clinical trials to treat cervical cancer (33). HDACi prevents HDACs from removing acetyl groups from proteins and this may stabilize NR2E3-enhanced p53 acetylation. We applied the ZIP drug synergy scoring tool (34,35) to assess the 11a-Romidepsin treatment combination, obtaining a ZIP average score of 10.5 and ZIP peak score of 25.4 (Fig. 5B). A ZIP average score > 10” suggests overall synergy between the drugs (34). A ZIP Peak score of 25.4 suggests the highest synergy and recommends the best dosage for each drug in a combination. Our data show that agonist 11a synergizes with Romidepsin and that the projected best dosage combination falls into their low dosage ranges. Low drug doses are favored for cancer patients because of the toxicities of anti-cancer therapy. One combination of 11a and Romidepsin from the low dosage range induced more cell apoptosis in one day than the individual single drug treatments (Fig. 5C). In fact, 11a or Romidepsin inhibited cell growth, whereas the 11a-Romidepsin combination killed most cells by 2 days (Fig. 5D).

**Figure 5:**
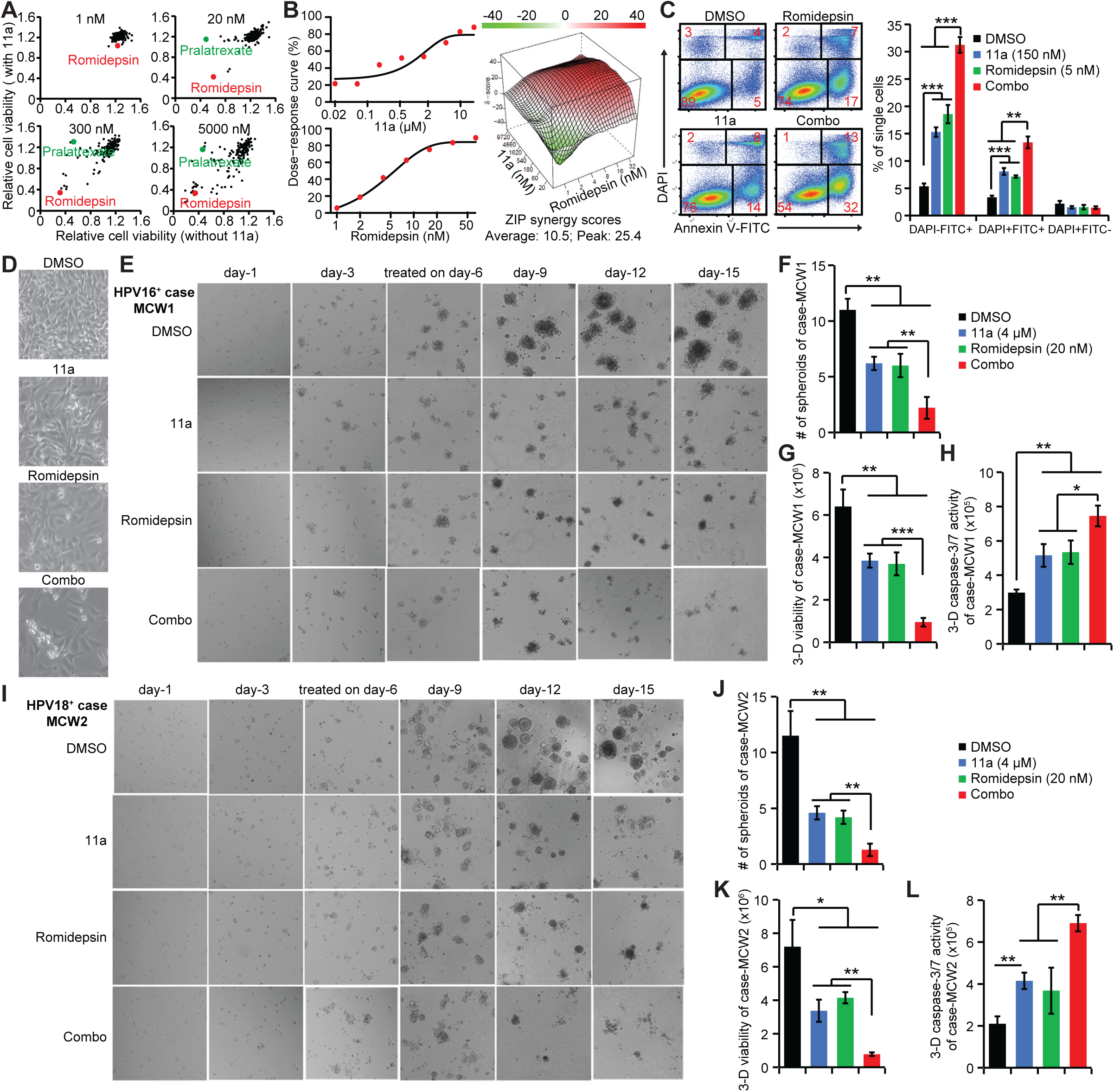
11a synergizes with Romidepsin to suppress cervical cancer. (A) Repurposing screens of FDA-approved anti-cancer drug library (AOD X) presented Romidepsin as a lead to promote 11a’s cell-killing effect. 4 concentrations of the drugs in the library were tested. The cell viability in DMSO control in each plate was normalized to 1. The normalized cell viabilities in the presence of 11a were plotted against those in the absence of 11a per concentration. (B) ZIP drug synergy score showed the synergy between 11a and Romidepsin. 8x9 doses of 11a and Romidepsin were mixed with HeLa cells in a 96-well plate. The cell viability in DMSO control was normalized to 100%. The data were evaluated for synergy by ZIP score(34). (C) The 11a-Romidepsin combo induced more HeLa cell apoptosis than each individual drug in 1 day in Annexin V-binding assay. (D) The combo had much stronger inhibition of 2-D HeLa cell growth than each individual drug in 2 days. (E-L) 3-D spheroid culture of two patient-derived cervical cancer cells MCW1 (E-H) and MCW2 (I-L). (E) The combo eliminated spheroids from patient-derived cervical cancer cells MCW1. The treatment started on day-6 when the spheroids were thriving. (F) The combo markedly decreased the number of MCW1 spheroids on day-15. (G) The combo markedly suppressed the spheroid viability of MCW1 in CellTiter-Glo 3-D assay on day-15. (H) The combo stimulated the caspase 3/7 activity in MCW1 spheroids in Caspase-Glo 3/7 3D Assay on day-15. (I) The combo eliminated spheroids from patient-derived cervical cancer cells MCW2. (J) The combo markedly decreased the number of MCW2 spheroids on day-15. (K) The combo markedly suppressed the spheroid viability of MCW2 on day-15. (L) The combo stimulated the caspase 3/7 activity in MCW2 spheroids on day-15. *p* value was calculated by Student’s t-test with two tails. *: *p*<0.05; **: *p*<0.01; ***: *p*<0.001.

3-D spheroid cultures can help predict the efficacy of 11a-Romidepsin treatment *in vivo* because they better represent the cervical cancer microenvironment and drug infiltration than the 2-D culture. 3-D spheroid cultures of HPV^+^ cervical cancer cell lines HeLa and SiHa confirmed the synergy between 11a and Romidepsin in suppressing spheroid size and quantity, lowering spheroid viability and inducing spheroid apoptosis by downstream activation of caspase-3/7 via p53 (36) (supplementary Fig. S8). To create additional spheroids consisting of more heterogeneous cell populations that are more representative of patient cancers, we employed two primary cervical cancer cell cultures MCW1 (HPV16^+^) and MCW2 (HPV18^+^). As shown in Figure 5E, patient-derived MCW1 cells started to form spheroids on day-3 and continued to thrive on day-15. The addition of 11a or Romidepsin on day-6 slowed the growth of the spheroids, while the combination of 11a and Romidepsin began to eliminate the spheroids on day-9 (Fig. 5E-G). Caspase 3/7 activities were significantly increased by these drugs, alone or in combination (Fig. 5H). We observed nearly identical results for spheroids grown from patient-derived MCW2 cells (Fig. 5I-L). Together, our data demonstrate the synergy between 11a and Romidepsin in suppressing the growth of human cervical cancer cells with superior translational values.

### Multifaceted anti-cancer mechanisms of 11a-Romidepsin combo

We first studied 11a’s specificity for NR2E3. The dose-dependent activation of the p53 reporter by 11a in the cells enforcedly expressing NR2E3 was stronger than in these enforcedly expressing NR2E1, NR2F1 and NR2F2 (supplementary Fig. S9A), suggesting a higher specificity of 11a for NR2E3. 11a’s specificity for NR2E3 was reaffirmed by knocking down NR2E3 in the p53 reporter assay (supplementary Fig. S9B). We next examined transcriptomics changes that were induced by 11a in HeLa cells (supplementary Fig. S10A-B) and confirmed the transcription changes of 6 cancer-associated genes by qRT-PCR (supplementary Fig. S10C). Interestingly, 11a repressed the transcription of *ABCB1* which encodes the Multidrug Resistance Protein, a key ATP-dependent drug efflux pump for anti-cancer drugs and is responsible for drug resistance in many cancer types (37). This finding raises the possibility that 11a may help counteract drug resistance in cancer cells. We also detected up-regulation of many pro-apoptosis genes, among which *CHAC1* is downstream of *ATF3* and increased by *NR2E3* (38). Consistent with the activation of NR2E3, 11a stimulated the p53, apoptosis and TGFβ gene sets while suppressing the oncogenic E2F and PI3K cancer pathways (supplementary Fig. S10D-F). 11a also suppressed ATP production through inhibiting both Glycolysis and Oxidative phosphorylation (supplementary Fig. S10G) in a manner similar to that caused by enforced expression of *NR2E3*. Treated HeLa cancer cells also activated the KRAS and TNFα pathways (supplementary Fig. S10H), which could represent alternate survival signals to help counteract the growth inhibitory actions of 11a on HeLa cells.

An RNA sequencing study with the 11a-Romidepsin combination displayed well-separated sample clusters, supporting synergy between 11a and Romidepsin (Fig. 6A-C). Up-regulation of six tumor-suppressing genes were verified by qRT-PCR (Fig. 6D). Of note, Romidepsin increased transcription of *ABCB1* in HeLa cells, but this was abolished by 11a (Fig. 6E). Romidepsin has been shown to up-regulate *ABCB1* in several cancer types, forming the basis for Romidepsin resistance in cancer patients (39,40). This resistance could be reversed by 11a.

**Figure 6:**
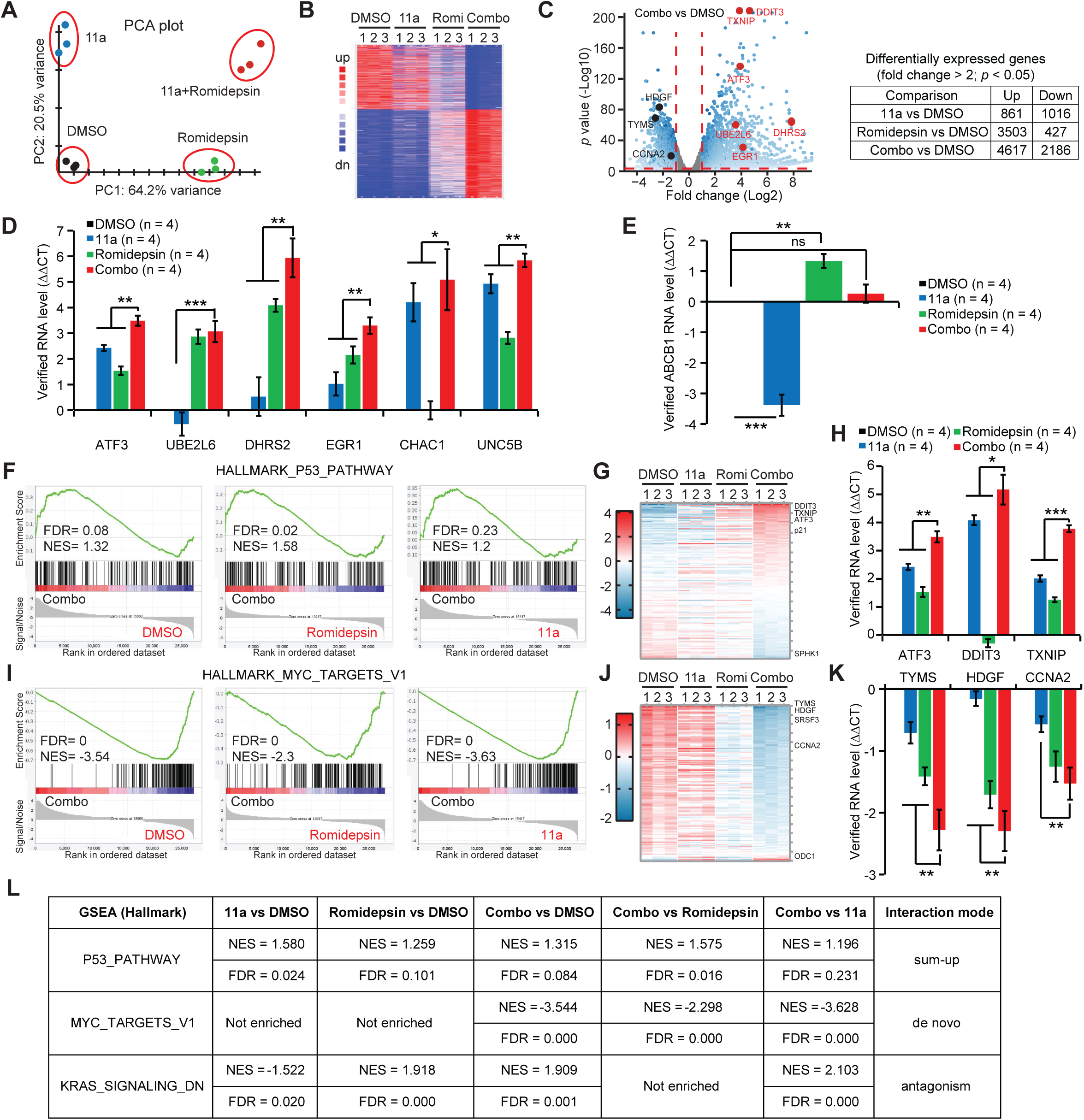
Dissecting molecular mechanisms underlying the drug synergy. RNA sequencing analysis of HeLa cells treated with the indicated drugs. (A) PCA plot indicated 4 well-separated sample clusters. (B) Heat map of differentially expressed genes in 4 groups. (C) Volcano plot of the transcripts of the combo vs DMSO. *Red*: growth inhibitory genes and *black*: growth promoting genes. The table summarized the number of differentially expressed genes in each comparison. (D) Up-regulations of 6 growth inhibitory genes were verified using qRT-PCR. (E) 11a reversed Romidepsin-induced up-regulation of Multidrug Resistance Protein ABCB1. (F) GSEA plots of p53 pathway in the indicated comparison. (G) Heat map of differentially expressed genes in the p53 pathway among 4 groups. (H) Up-regulations of p53 pathway genes were verified using qRT-PCR. (I) GSEA plots of MYC pathway in the indicated comparison. (J) Heat map of differentially expressed genes in the MYC pathway among 4 groups. (K) Down-regulations of MYC pathway genes were verified using qRT-PCR. (L) Three drug synergy modes were identified by multiple comparisons of GSEA. *p* value was calculated by Student’s t-test with two tails. *: *p*<0.05; **: *p*<0.01; ***: *p*<0.001.

Multiple comparisons of GSEA between DMSO, 11a, Romidepsin and the 11a-Romidepsin combination were conducted to further understand these drug synergies. Some pathways such as p53 were activated by 11a and Romidepsin, respectively, and were further activated by the drug combination (Fig. 6F). Pro-apoptotic genes (*e.g., DDIT3*, *TXNIP* and *ATF3*) in the p53 pathway were up-regulated most by the drug combination, which was verified by qRT-PCR (Fig. 6G-H). We named this drug synergy the “sum-up” mode (Fig. 6L). In contrast, the MYC oncogenic pathway was not inhibited by either 11a or Romidepsin alone but was repressed by the 11a-Romidepsin combination (Fig. 6I). Growth-promoting genes (*e.g., TYMS*, *HDGF* and *CCNA2*) in the MYC pathway were down-regulated most by the drug combination (Fig. 6J-K). We named this type of drug synergy the “*de novo*” mode which was only seen in the combo (Fig. 6L). A third type of synergy was the “antagonism” mode in which survival signals activated by either 11a or Romidepsin were antagonized by the other to cause cell death. For example, 11a down-regulated the KRAS-down pathway while Romidepsin up-regulated these genes (Fig. 6L). The 11a-Romidepsin combination up-regulated the KRAS-down genes, suggesting that Romidepsin antagonized and overcame 11a’s effects on activating the KRAS-down pathway (Fig. 6L). We observed additional examples of these synergy modes (supplementary Fig. S11). Together, our data revealed the multifaced molecular synergies of 11a and Romidepsin in suppressing cervical cancer cells.

## Discussion

### Mutations and short isoform of *NR2E3* lack anti-tumor functions

We aim to address the tumor-suppressing functions of *NR2E3*. We have continued increasing our knowledge of NR2E3’s p53-dependent functions by showing its activation of wild-type and mutated p53 in multiple cancer cell lines and explored its p53-independent functions by identifying the up-regulation of IFNα pathway and the down-regulation of MYC pathway (Fig. 1). Secondly, we discovered the higher frequencies of *NR2E3* nonsynonymous SNVs in cancer patients than in regular population (Fig. 2), which encouraged us to use the p53 reporter as the tool to stratify pathogenic *NR2E3* SNVs such as *NR2E3^R97H^*(Fig. 3). R97H that was detected in uterus cancer lacks all of the p53-dependent and -independent anti-tumor functions of wild-type *NR2E3* (Fig. 4). Surprisingly, the short isoform of NR2E3 from alternative RNA splicing acts as dominant-negatively (Fig. 1). Together, our studies of both wild-type and mutated *NR2E3* indicate NR2E3 as a tumor suppressor.

That NR2E3 is a tumor suppressor has shed light on *NR2E3*-associated diseases. First, there are observations of a later-stage dysplastic appearance and a proliferative response only in human retinal degenerations bearing mutated *NR2E3* (13). Our studies suggest that mutated *NR2E3* loses the capability to induce cell apoptosis, leading to dysplasia and overgrowth of human retinal cells. Secondly, our studies have brought attentions to the *NR2E3* SNVs detected in cancer cases. Of note, these reported *NR2E3* SNVs are all germline. Do they play roles in cancer predisposition?

Though higher than in other tissues, *NR2E3* expression is still quite low in the urogenital system, which has proven challenging for us to study *NR2E3* SNVs at physiological levels using tools such as Crispr/Cas9 to knock-in SNVs under control of the endogenous *NR2E3* promoter. Even so, our enforced-expression assays showed reproducible differential effects on p53 transactivity, apoptosis, p53 acetylation and other cell growth processes by wild-type versus natural and engineered *NR2E3* and *p53* mutants. Significantly, the NR2E3 agonist 11a also exhibited expected p53, apoptosis and growth inhibitory effects on cervical cancer cell lines and 3D spheroids from cervical cancer patients (*see discussion below*). These congruent experiments support the physiologic validity of our data. Therefore, the involvement of *NR2E3* SNVs in cancer can be studied in transgenic mouse models of *NR2E3* (24). For instance, *NR2E3^R97H^* and *SP100^H727R^*, a tumor suppressor (41), co-exist in one TCGA uterus endometrial epithelium cancer case. Their oncogenic roles can be studied with conditional expression of *SP100^H727R^* in uterus using *BAC-Sprr2f-Cre* (42) in germline *NR2E3^R97H^* knock-in mouse model.

### Therapeutical potential of the NR2E3 agonist 11a

We previously reported the wide-spectrum of anti-cancer roles of 11a in NCI-60 cancer cell panel (22). Currently, the results of 3-D spheroid culturing patient-derived cervical cancer cells (HPV16^+^ and HPV18^+^) consisting of mixed cell populations indicate that 11a is able to infiltrate and kill the tumor cells in the presence of Romidepsin, highlighting its value in solid tumors (Fig. 5 & supplementary Fig. S8). In addition to cervical cancer, 11a synergizes with Romidepsin, Bortezomib or Carfilzomib to kill multiple myeloma cells and retinoblastoma cells which endogenously express wild-type p53 and NR2E3 (*data not shown*). More interestingly, 11a synergizes with a FOXM1-inhibitor to kill multiple myeloma cells bearing p53^R175H^ mutation (*see discussion below*). Undoubtedly, NR2E3 agonist shows its therapeutical values in cancer alone and in combination.

11a preferentially stimulates endogenous NR2E3 and p53 (supplementary Fig. S7-10). We are interested in exploring the molecular functions of 11a. For example, 11a suppresses *ABCB1* which is a major cause of resistance to many anti-cancer drugs, including Romidepsin (37,39,40). The up-regulation of *ABCB1* by Romidepsin is abolished by 11a (Fig. 6E), highlighting the value of 11a in combinatorial therapies. The growing knowledge of 11a’s molecular functions also helped to computationally identify the synergy between 11a and a recently-invented FOXM1 inhibitor NB73 (43,44) by aligning the RNA sequencing data between 11a-treated HeLa cells and *FOXM1*-deleted OPM2 multiple myeloma cells (*data not shown*). The predicted synergy in OPM2 cells which endogenously express p53^R175H^ mutation was experimentally verified (*data not shown*). Collectively, it seems clear that NR2E3’s agonist 11a deserves further investigation, including pharmacokinetics and pharmacodynamics studies in preclinical models, with a continued eye to possible human cancer treatments.

## Supporting information

supplemental will support the main text

## Acknowledgement

We deeply thank Dr. Paul Ahlquist and Dr. Wei Xu at Institute of Molecular Virology, McArdle Laboratory for Cancer Research, University of Wisconsin-Madison who provided mentorship and numerous supports of materials and team members to this project. We also thank Drs. Bill Sugden, Shiming Chen, Arthur Polans, Yeping Sun and Bert Vogelstein for generously sharing reagents and materials and scientific discussion. Lance Rodenkirch at Optical Imaging core facility, University of Wisconsin-Madison supported this project.

This work was supported by the Tom and Sally Ebenreiter Precision Medicine Research Award and Marshfield Clinic Research Foundation startup fund to Z.W. and by the NIH through grants CA-22443, CA-125387, CA-014520, CA-151354, CA-262198 and OD-026555 to P.A. (supporting L.H.), W.X. (supporting Y.W.), S.H., J.S.R., S.J., *All of Us*-Wisconsin (S.H.) and UW Comprehensive Cancer Center (UWCCC).

## Authorship

Contribution: Z.W. conceived, designed and supervised the study; Y.W., T.K., L.H., V.Z. and Z.W. designed and conducted the experiments and interpreted the data; J.M., Z.W. and S.H. conducted the NR2E3 mutation-cancer association study; J.S.R. established the patient-derived cervical cancer cell strains; A.S., D.P., and M.I. completed the 3-D spheroid experiments; L.M., S.K.S., S.G. and A.B. provided technical or material support; and Y.W., T.K., G.A., D.P., J.S.R., S.J. and Z.W. wrote, reviewed, and/or revised the manuscript.

Conflict-of-interest disclosure: The authors declare no competing financial interests.

## Data Assess Statement

We will follow the policies of NIH and Cancer Discovery journal to share the data.

## Figure legends

**Figure S1.**
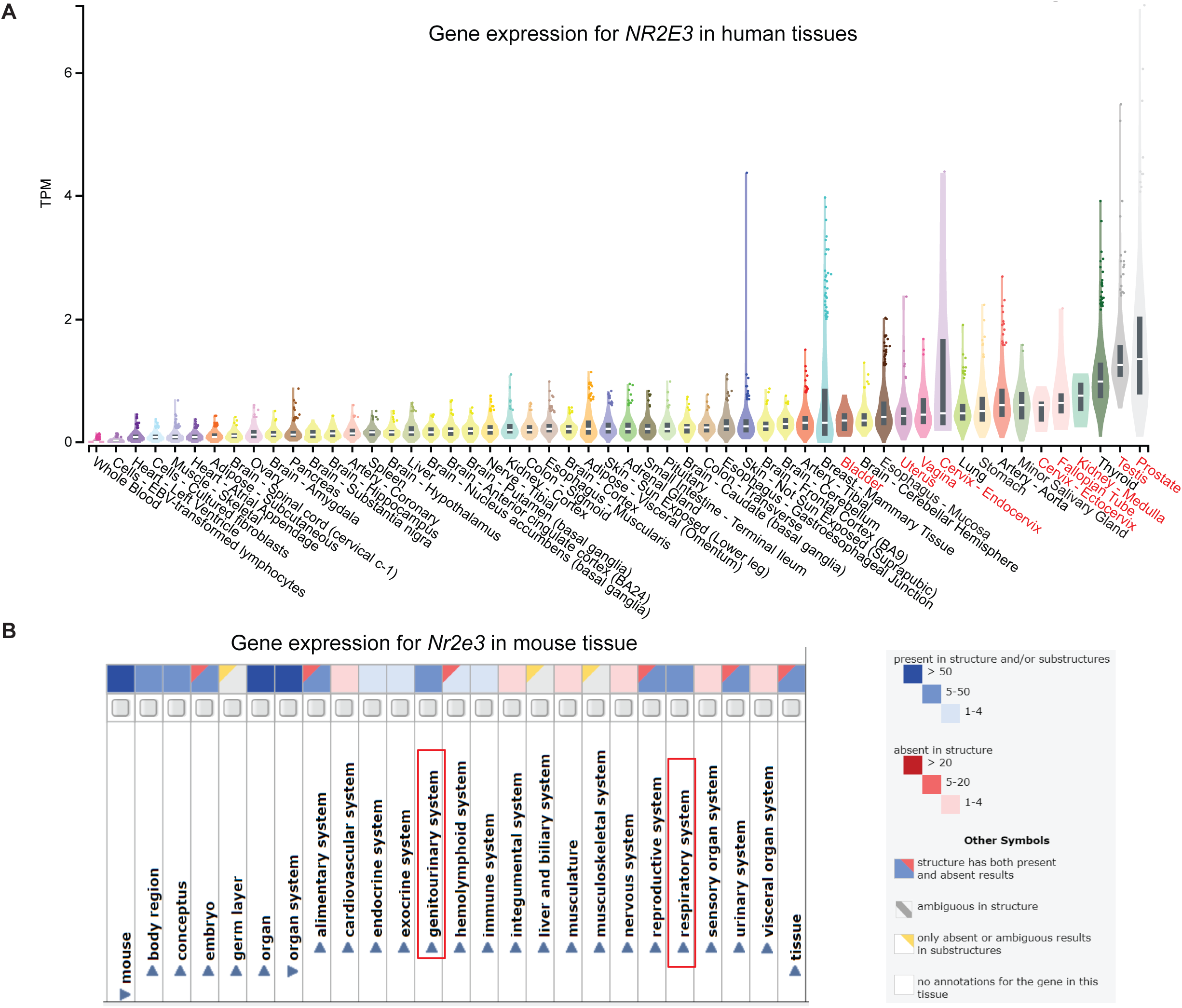

**Figure S2.**
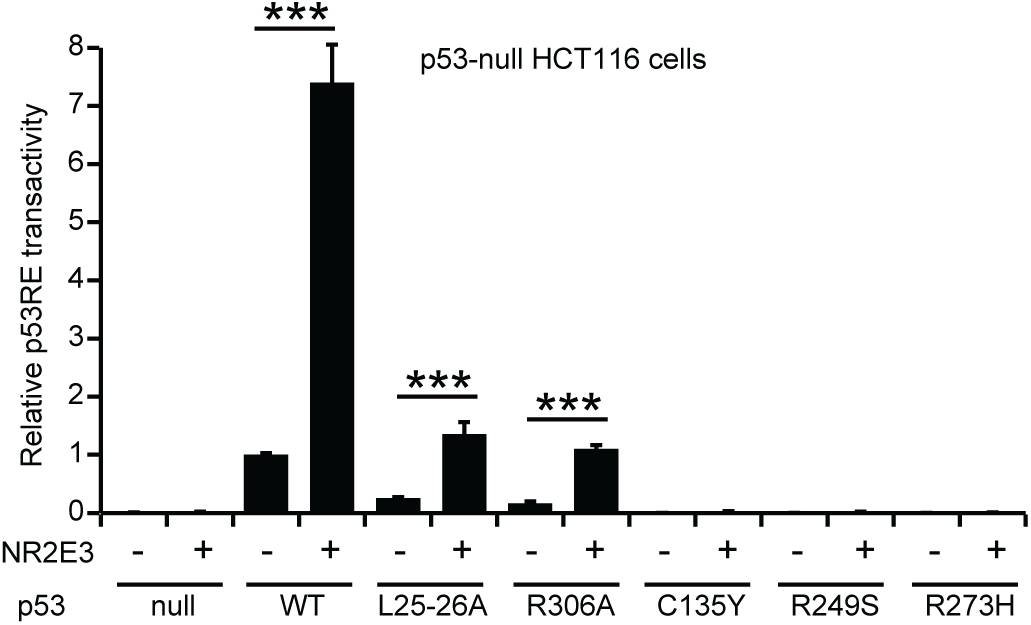

**Figure S3.**
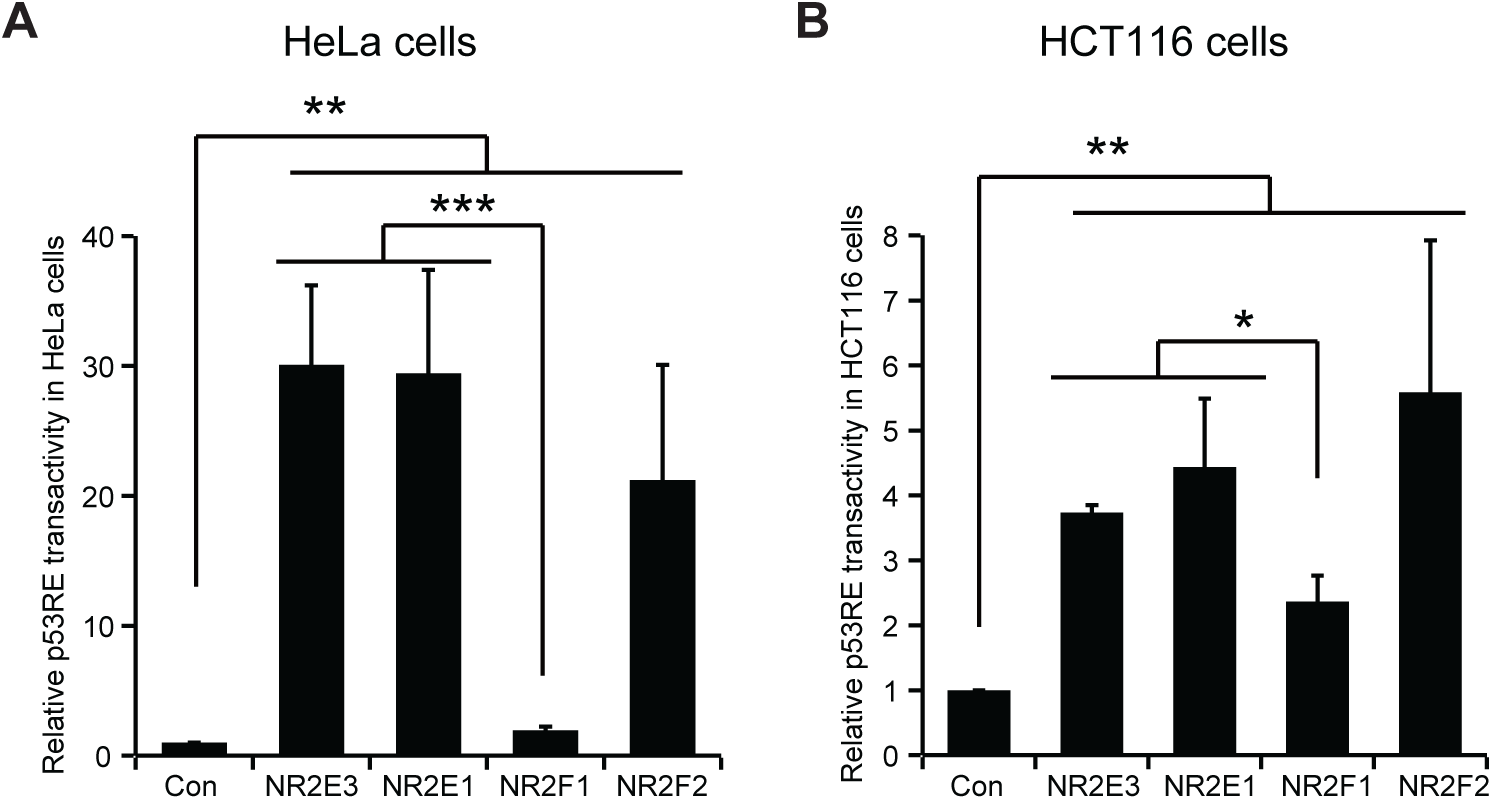

**Figure S4.**
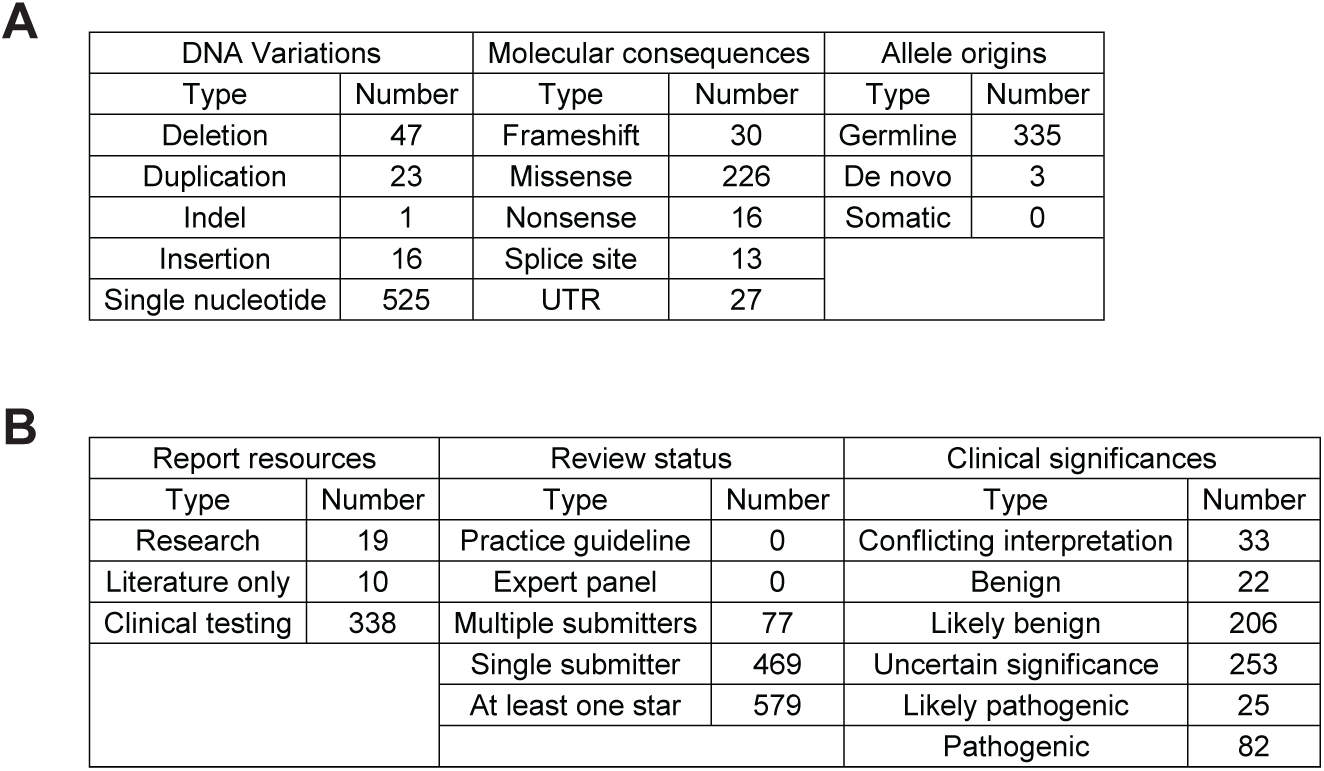

**Figure S5.**
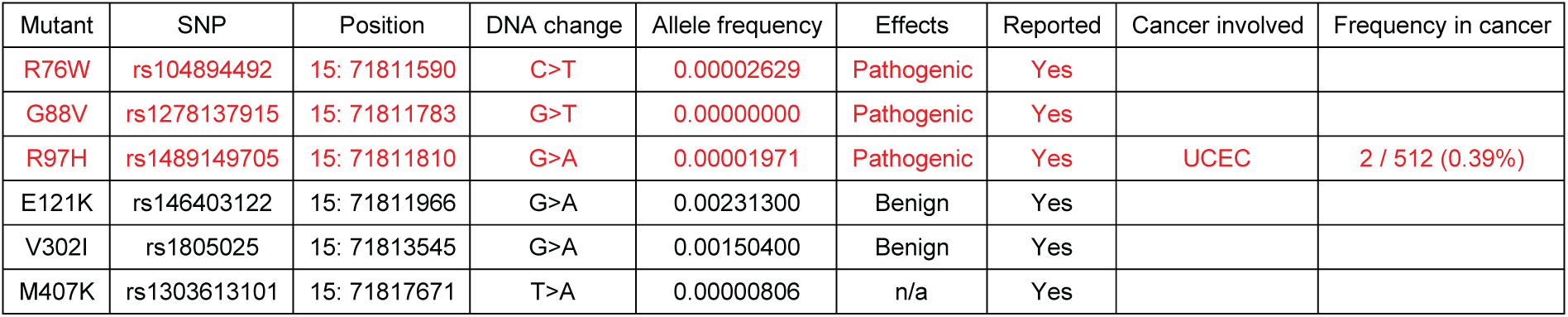

**Figure S6.**
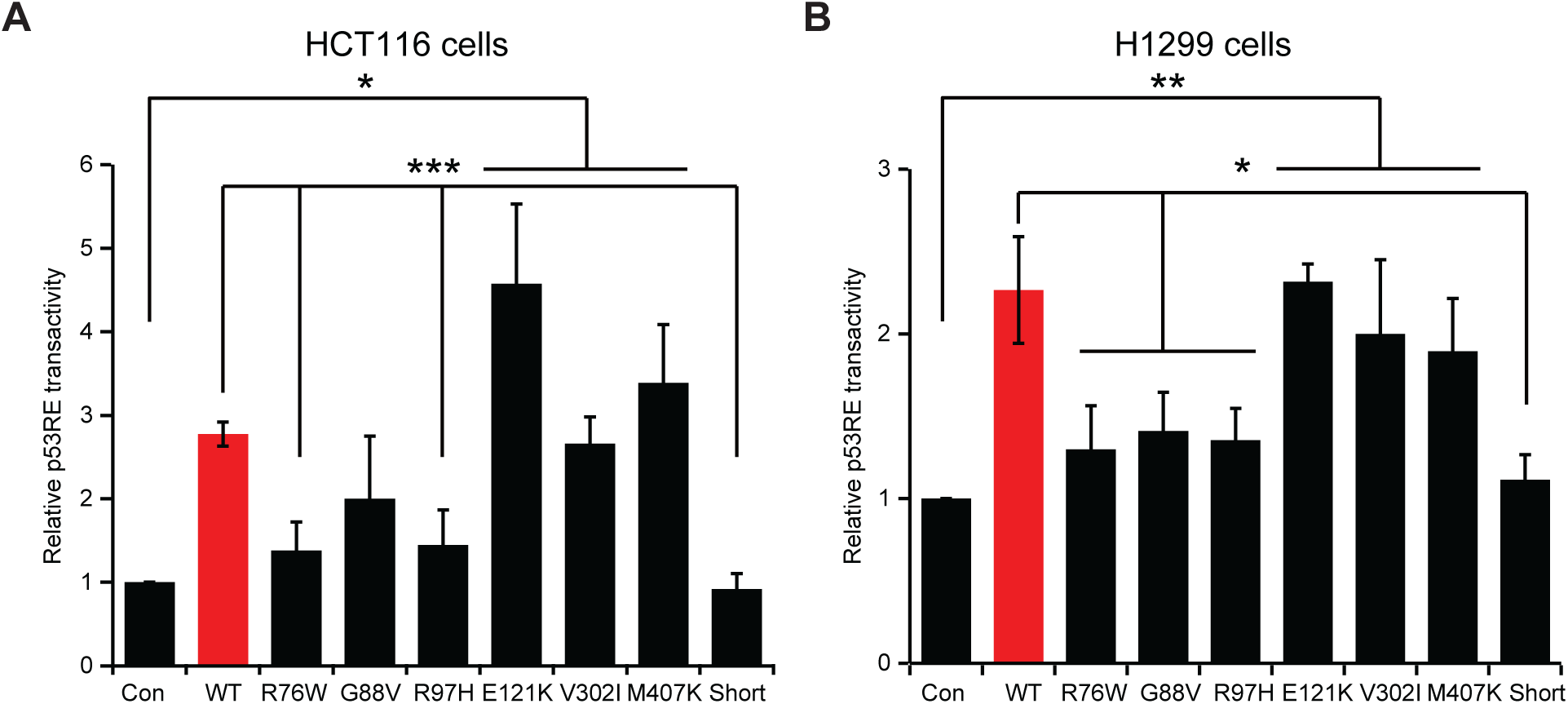

**Figure S7.**
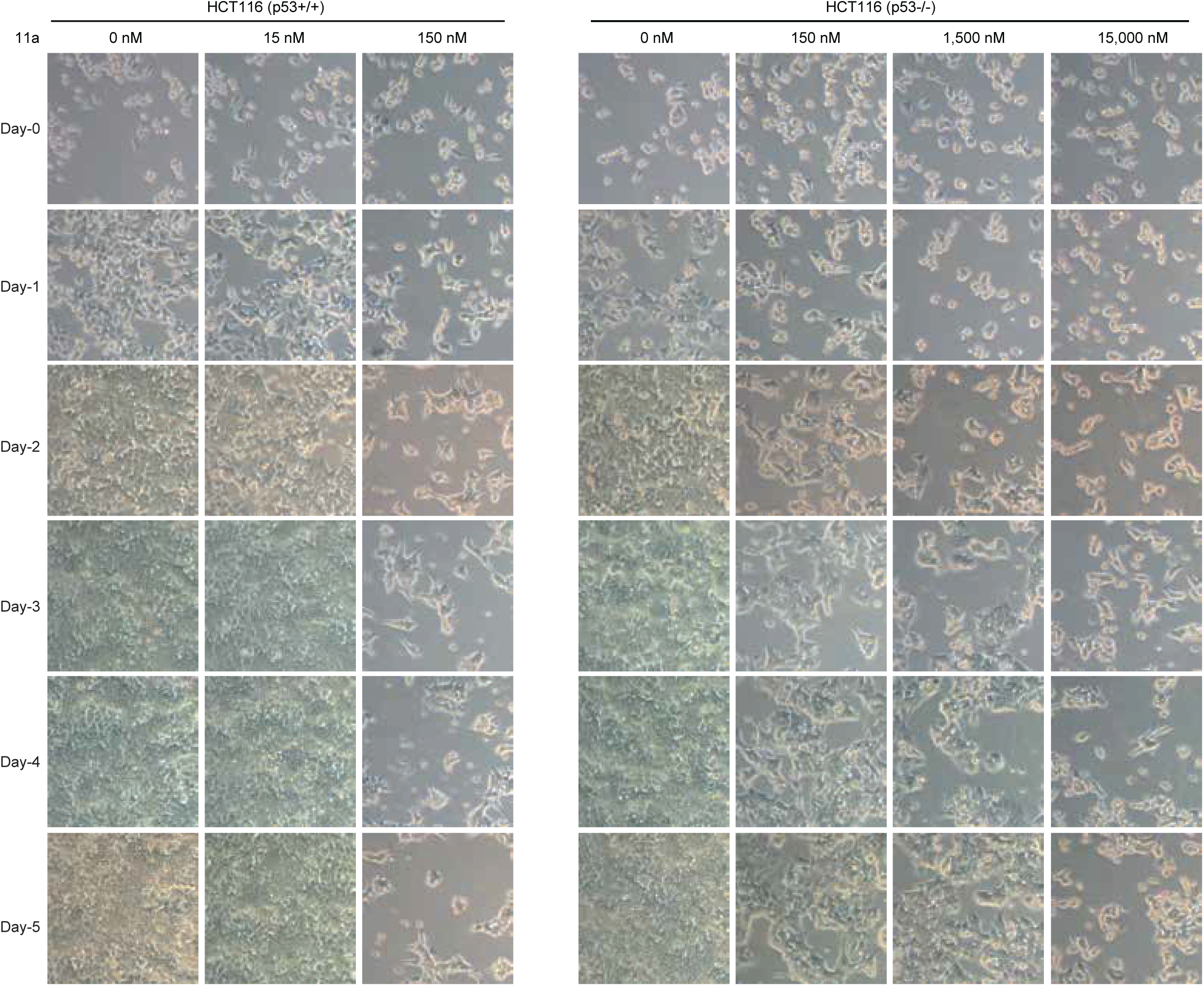

**Figure S8.**
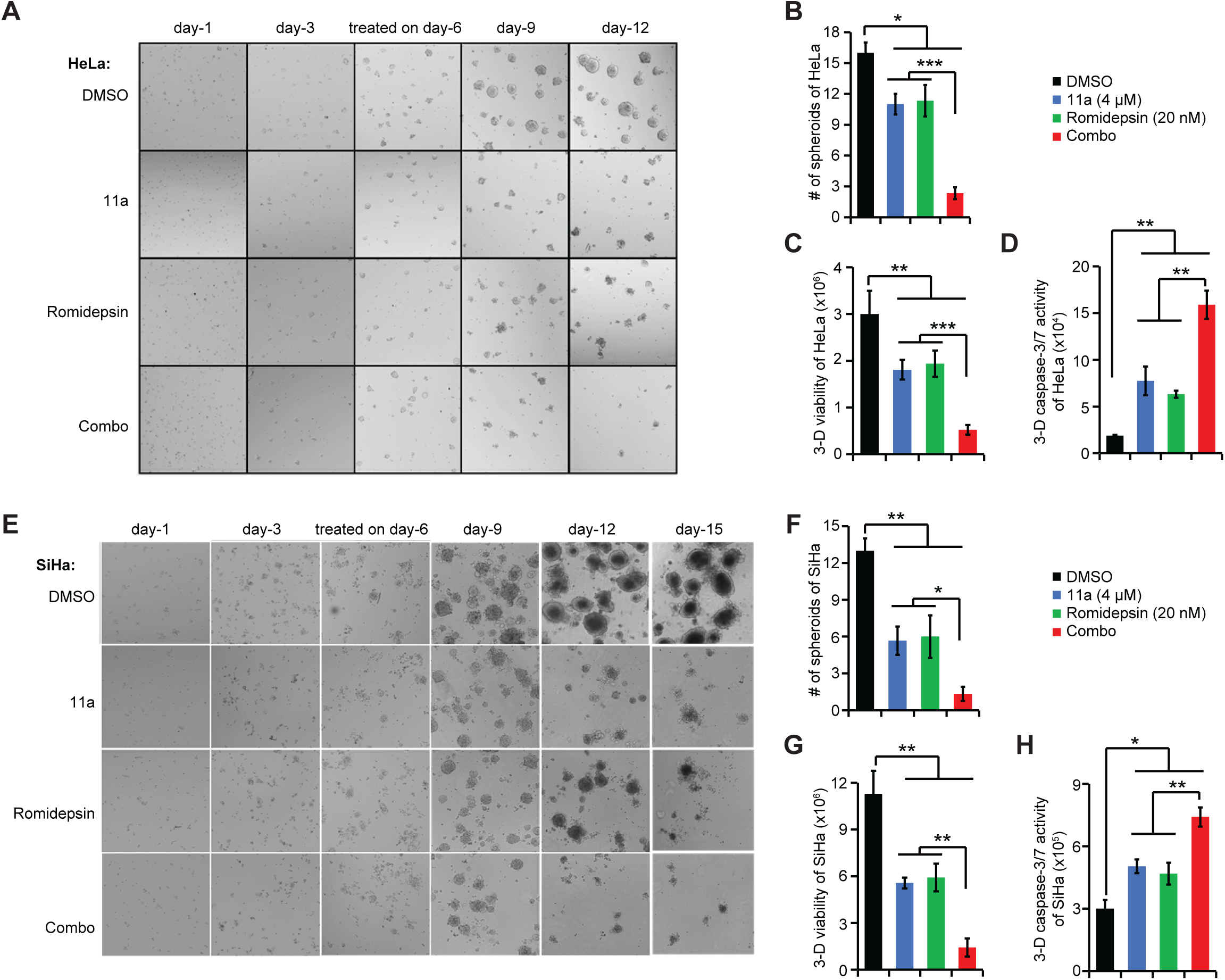

**Figure S9.**
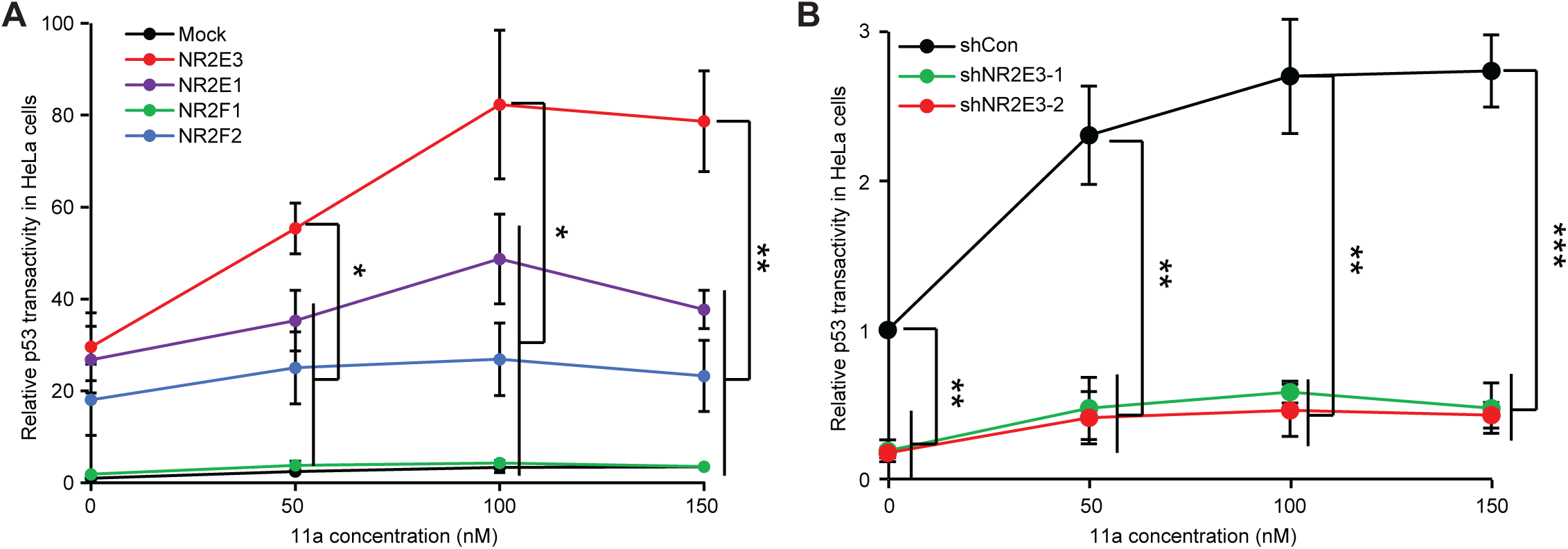

**Figure S10.**
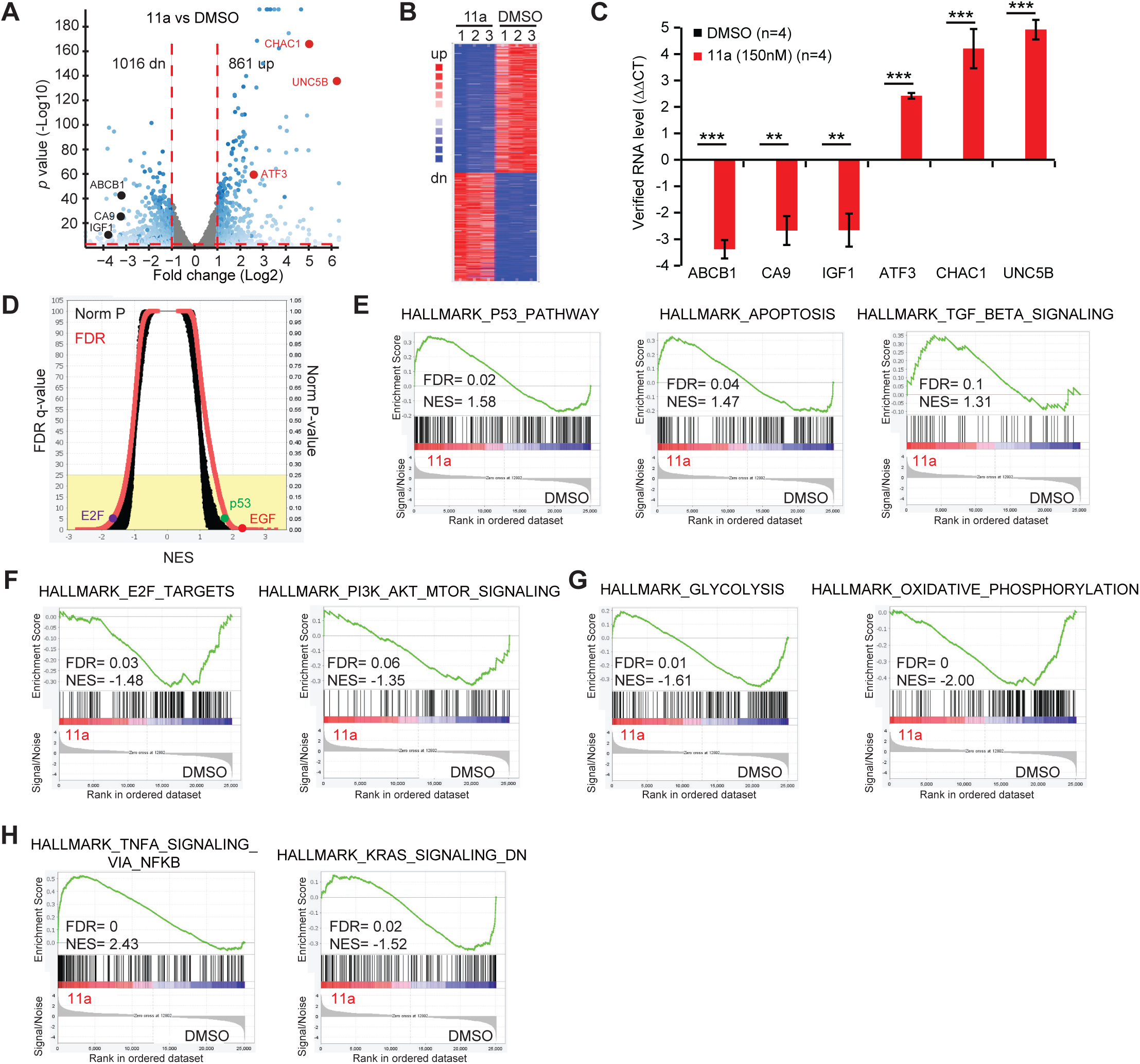

**Figure S11.**
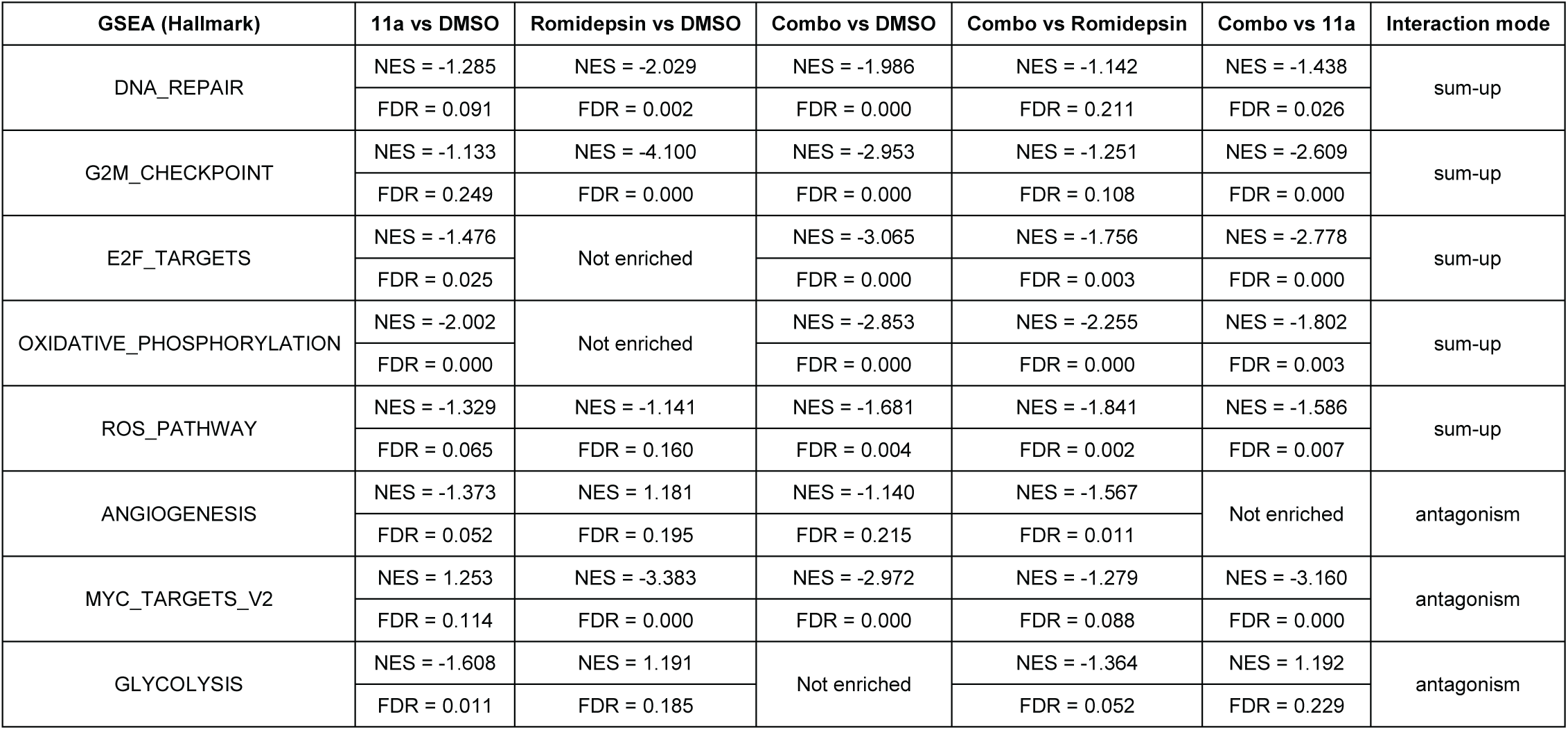

