## supplemental will support the main text for "Orphan nuclear receptor *NR2E3* and its small-molecule agonist induce cancer cell apoptosis through regulating p53, IFNα and MYC pathways"

**Running title:** Activating *NR2E3* suppresses cancer
**Key words:** Orphan nuclear receptor *NR2E3*, loss-of-function mutations, anti-cancer drug synergy, 3D spheroid culture, tumor suppressor p53

### Materials and Methods:

**Plasmids and cell lines.** Human NR2E3 full-length, p53-C135Y, p53RE-SEAP, p53RE-FLuc, and SV40-RLuc were described (1). All NR2E3 and p53 mutations were achieved with QuickChange II XL site-directed mutagenesis kit (Agilent, cat# 200521) followed by Sanger sequencing. NR2E3 truncates were made by PCR with indicated primer sets.

Y79, HeLa, SiHa and MCF7 cells were purchased from ATCC. *p53*$^{-/-}$ and *p53*$^{+/+}$ HCT116 cells were kindly provided by Dr. B. Vogelstein, John Hopkins University, H1299 cells by Dr. W. Sugden, University of Wisconsin-Madison, RKO cells by Dr. P. Lambert, University of Wisconsin-Madison, and MM1S and OPM2 cells by Dr. S. Janz, Medical College of Wisconsin. FBS (GIBCO, cat# 16000069) was heat-inactivated at 56$^{o}$C for 30 minutes and added to a final 10% concentration into DMEM (GIBCO, cat# 11-140-050) for HeLa, SiHa, H1299, RKO and MCF7 cells, RPMI-1640 (ATCC, cat# 50-188-266FP) for MM1S and OPM2 cells, and McCoys'5A (Hyclone, cat# SH30200.FS) for HCT116 cells, and to a final 20% concentration into DMEM for Y79 cells. All the cell lines were cultured at 37$^{o}$C in cell incubators filled with 5% $CO_2$ and passaged every two or three days.

**Repurposing screens of FDA-approved anti-cancer drug library.** The AOD X library was kindly provided by Developmental Therapeutics Program, Division of Cancer Treatment and Diagnosis, National Cancer Institute. This library contains 166 FDA-approved anti-cancer drugs at 10 mM in 20 µL DMSO. These drugs were re-plated into 384-well Echo-compatible source plates (Beckman-Coulter, cat# C74290). Echo Plate reformat software was used to set up the cross-titration matrix of drugs. Drug combinations were spotted into white TC-treated clear bottom 384-well assay plates (Corning, cat# 3765) using the Beckman-coulter Echo 650 acoustic liquid handler. All assay points were backfilled with DMSO to a final volume of 50 nL per well. The assay plates were sealed with sterile plate seals and frozen at -20$^{o}$C until needed.

50 µL $0.8x10^5$/mL HeLa cells or $3x10^5$/mL Y79 cells were seeded to each well of the assay plates by BioTek MultiFlo microplate dispenser with 5-µL stainless steel cassettes. The final concentration of 11a was 1.5 µM for HeLa cells and 4 µM for Y79 cells, respectively. After brief centrifugation at 200x g, the cells were cultured at 5% $CO_2$, $37^oC$ for two days. 20 µL CellTiter-Glo cell viability substrate (Promega, cat# G7573) were added per well with BioTek MultiFlo microplate dispenser with 5-µL plastic cassettes and incubated at room temperature for 10 minutes on a microplate shaker (USA Scientific, cat# 7402-4000). The luminescence was measured with BMG LUMIstar Omega plate reader. The cell viability in the DMSO control in each plate was normalized to 1. The normalized cell viabilities in the presence of 11a were plotted against those in the absence of 11a for each assayed concentration of the AOD X library. The outliers were selected by naked eyes.

**ZIP drug synergy scoring assay.** Romidepsin, Bortezomib and Carfilzomib (10 mM in DMSO) were purchased from TargetMol (Wellesley Hills, MA). 11a was synthesized by Pharmabridge Inc. (Doylestown, PA) and dissolved to 10 mM in ethanol alcohol and then DMSO (2). NB73 was kindly provided by J. Katzenellenbogen (3) (University of Illinois at UC) and dissolved to 5 mM in ethanol alcohol with sonication and then DMSO.

The drugs were sequentially diluted in Opti-MEM medium (GIBCO, cat#31985070) in 8-stripe PCR tubes. 5 µL each diluted drug was aliquoted with 8-channel P20 pipetman (Gilson, cat# F144070) to each well in a 96-well plate. 100 µL $1.2x10^5$/mL HeLa/MCF7 cells or $3x10^5$/mL Y79/MM1S/OPM2 cells were seeded to each well with 8-channel P200 pipetman (Gilson, cat# F144072). The cells were cultured at 5% $CO_2$, $37^oC$ for two days. 20 µL CellTiter-Glo cell viability substrate (Promega, cat# G7573) were added per well with DISTRIMAN Repetitive Pipette (Gilson, cat# F164001) and incubated at room temperature for 10 minutes on the microplate shaker (USA Scientific, cat# 7402-4000). The luminescence was measured with BMG LUMIstar Omega plate reader.

The cell viability in the DMSO control was normalized to 100%. ZIP synergy score was used to evaluate the synergy, following the instructions at https://synergyfinder.fimm.fi (4). "ZIP Average Score > 10" suggests a synergy between drugs; "ZIP Average Score < -10" suggests an antagonism between drugs; "ZIP Average Score between -10 and 10" suggests an additive effect between drugs.

**3-D spheroid culture of patient-derived cell lines and ATCC cell lines of cervical cancer.** We established two cervical cancer patient-derived cell lines MCW1 that is $HPV16^+$ and MCW2 that is $HPV18^+$. They were grown from patient tumor explants in serum-free media supplemented with KGM. The cells have passed ATCC STR test as mycoplasma negative. All operations and experiments were reviewed and approved by the MCW institutional review board. MCW1 and MCW2 cell lines were maintained in F-Media with 10% fetal bovine serum (GIBCO, cat# 16000069) and 1x penicillin-streptomycin (GIBCO, cat# 10-378-016) while ATCC cervical cancer cell lines HeLa and SiHa were maintained in DMEM (GIBCO, cat# 11-140-050). They were all cultured at $37^oC$ in cell incubators filled with 5% $CO_2$. All cells were negative for mycoplasma contamination.

*Generation of 3-D spheroids:* Cervical cancer cells were cultured as 3-D spheroids as described previously (5-7). Briefly, MCW1, MCW2, HeLa and SiHa cells were seeded at a density of 7,000 cells in Clonal Cells TCS Methylcellulose Medium (Stemcell Technologies, cat# 03814) mixed with DMEM at 1:1 ratio in 24 wells Ultra-low attachment plate (Sigma, cat# CLS3473). Cells were incubated at $37^oC$ in cell incubators filled with 5% $CO_2$ for 15 days and photographed of spheroids on day-0, 3, 6, 9, 12 and 15 with Nikon (Eclipse Ts2R) Microscope under 20X magnification. Media was changed after every 2 days initially (first 2 to 4 days) then every day by carefully aspirating 50% of the well volume and replacing with fresh complete media (Clonal Cells media and DMEM at 1:1). 11a, Romidepsin and their combo were added at the indicated concentrations on day-6. The experiment was terminated on day-15. 3-D spheroids were counted under microscope in each quadrant on day-15.

*3-D spheroids viability assay and apoptosis assay:* 3-D spheroids viability assay and apoptosis assay were performed as described previously (8-10). In brief, 2,000 cells of MCW1, MCW2, HeLa and SiHa were seeded per well in 96-well U-Bottom plate (faCellitate, cat# F202003) in triplicates with 100 µL of Clonal Cells media and DMEM (1:1), respectively. 48 hours later, old media was replaced with fresh Clonal Cells media containing the indicated drugs. 3-D spheroid cells viability and caspase-3/7 activity were analyzed after 24 hours using CellTiter-Glo 3D Cell Viability Assay (Promega, cat# G9681) and Caspase-Glo 3/7 3D Assay (Promega, cat# G8981) with Molecular Devices SpectraMax bioluminescent microplate reader.

**Bulk RNA sequencing.** To study NR2E3 wild-type and R97H, 2 µg of each plasmid (supplemented with empty vector to a total 4 µg) were transfected into HeLa cells with 8 µL TransIT-LT1 (Mirus, cat# MIR2300) 18 hours after $1x10^6$ HeLa cells were seeded in 6-cm dishes. 1 day after transfection, the medium was replaced with fresh medium. 2 days after transfection, RNA was extracted with RNeasy Mini kit (Qiagen, cat# 74106). RNA quantity and quality assessment was done with Nanodrop and Bioanalyzer 2100, library construction of mRNA was completed with NEBNext Ultra II RNA Library Prep kit for Illumina (NEB, cat# E7765) following the manual instruction, quantity and size distribution of the library was evaluated with Qubit and Bioanalyzer 2100, and libraries were pooled at equal molar concentrations, loaded on a NovaSeq 6000 S4 flow cell, and run on a NovaSeq 6000 sequencer. Each sample was sequenced at 20M reads with 2x150bp paired end.

To study the drug combination, 150 nM 11a and/or 5 nM Romidepsin were added to HeLa cells 1 day after $1x10^6$ HeLa cells were seeded in 6-cm dishes. RNA was extracted 1 day after drug treatments. RNA quantity and quality assessment was done with Nanodrop and Bioanalyzer 2100, library construction of mRNA was down with BGI in-house kit, quantity and size distribution of the library was done with Qubit and Bioanalyzer 2100, equal amount of each library was pooled and loaded to DNBSEQ-T7 PLATFORM, and each sample was sequenced at 20M reads with 2x150bp paired end.

The fastq files were aligned to hg38 reference genome and counted with STAR app and the differential expression between samples were acquired with DESeq2 app of RNA-Seq pipeline at www.basepairtech.com. GSEA 4.1.0 application was used to conduct the pathway analysis with the v2023.1 human gene sets (11).

**Chromatin Immunoprecipitation (ChIP).** HeLa cells were co-transfected with 1.2 µg p53 plasmid and 2.4 µg NR2E3 WT plasmid/empty vector in 10-cm dishes. Two days later, the cells were processed following the protocol which included the sequences of the PCR primers of p53BS1 and p53BS2 on p21 promoter and p53BS on MDM2 intron. Primer sequences were provided separately. Detailed ChIP protocol is following:

1. Two days after transfection, HeLa cells were washed twice with 10 mL PBS
2. Add 270 µL formaldehyde (from 37% stock, final is 1%) into 10 mL PBS per 10-cm dish, mix immediately and incubate at room temperature (RT) for 30 minutes
3. Add 1.25 mL 1.25M glycine and incubate at RT for 5 minutes with gentle shaking

4. Place the dishes on ice and wash once with 10 mL cold PBS
5. Harvest the cells with scrapers into 10 mL cold PBS and centrifuge at 1,500x rpm at 4$^{o}$C for 5 minutes in a Beckman-Coulter GPR centrifuge
6. Wash cell pellets with 10 mL buffer I (0.25% TritonX100, 10mM EDTA, 0.5 mM EGTA, 10 mM HEPES) and II (200mM NaCl, 1mM EDTA, 0.5mM EGTA, 10mM HEPES, pH 6.5) sequentially
7. Resuspend the cells in 1 mL lysis buffer (1% SDS, 10mM EDTA, 50mM Tris, pH 8.1) with protease inhibitors
8. Sonicate the cells with microtip (10 seconds at 50% power input with 2 minutes rest on ice for 8 cycles. Do not get too cold, or the SDS will precipitate in the buffer) using a Branson Digital Sonifier 450 sonicator
9. Transfer the cell lysates into 1.5-mL microtubes and centrifuge at 15,000x rpm at 10$^{o}$C for 15 minutes
10. Dilute the supernatant 10 fold with cold dilution buffer (1% TritonX100, 2mM EDTA, 150mM NaCl, 20mM Tris, pH 8.1) with protease inhibitor cocktails in 15-mL tubes on ice. Leave 100 µL aliquot as input control
11. Incubate the diluted chromatin solution with 3 µL anti-acetylated p53 antibody (Upstate, cat# 06-758) on a nutator in cold room for 12 to 16 hours
12. Add 50 µL 50% slurry of protein A-sepharose 4B beads (washed in TE pH 8.0; blocked with ssDNA) and incubate on the nutator in cold room for 1 hour
13. Centrifuge at 1,500x rpm at 4 $^{o}$C for 2 minutes to pellet the beads and discard the supernatant
14. Resuspend the beads in 1 mL TSE+150mM NaCl buffer (0.1% SDS, 1% TritonX-100, 2mM EDTA, 20mM Tris, 150mM NaCl, pH 8.1) and wash the beads with a Fisher Genie 2 Vortex at setting 1 in cold room for 10 minutes.
15. Spin and wash the beads again with 1 mL TSE+500mM NaCl buffer (0.1% SDS, 1% TritonX-100, 2mM EDTA, 20mM Tris, 500mM NaCl, pH 8.1) and then with buffer III (0.25M LiCl, 1% NP-40, 1% deoxycholate, 1mM EDTA, 10mM Tris, pH 8.1)
16. Wash beads twice with 1 mL TE buffer and elute the immune complexes with 300 µL 1% SDS, 0.1M $NaHCO_3$ with Fisher Genie 2 Vortex at setting 6 at RT for 30 minutes
17. Incubate the elutions and the aliquots (the input control) at 65$^{o}$C for 4 to 6 hours
18. Purify DNA with PCR purification kit (Qiagen, cat# 28104)
19. Use GoTaq PCR kit (Promega, cat# M7660) to amplify interested fragments (50 $^{o}$C, 2 minutes; 95 $^{o}$C, 10 minutes; 29 cycles of 95 $^{o}$C, 15 seconds; 55 $^{o}$C, 30 seconds and 72 $^{o}$C, 30 seconds; 72 $^{o}$C, 7 minutes; and 4 $^{o}$C forever) for electrophoresis in 2% agarose gel containing ethidium bromide staining with a thin comb for images
20. Use TaqMan™ Fast Advanced Master Mix (Applied Biosystems, cat# 4444557) and ABI 7900HT device to quantify the ChIP products

**Gel shift assay (EMSA).** HeLa cells were co-transfected with 0.6 µg p53 plasmid and 0.6/1.2 µg NR2E3 WT, or 1.2 µg R76W/R97H plasmids, respectively. Two days later, nuclear extracts were harvested (1). The sequence of p53-binding consensus and the protocol were provided separately. Detailed EMSA protocol is following:

1. Anneal DNA primers:

| | |
|---|---|
| 10x NEB buffer 2 | 10 µL |
| Primer 1 (100uM, 1.1 µg/µL) | 1 µL |
| Primer 2 (100uM, 1.1 µg/µL) | 1 µL |
| $H_2O$ | 38 µL |

PCR program: 94 $^{o}$C, 2 minutes; 65 $^{o}$C, 10 minutes; 37 $^{o}$C, 10 minutes; and 4 $^{o}$C forever

2. Label probe with $^{32}$P-dCTP:

5xbuffer 3 µL
Annealed Probe 2.5 µL
Klenow (exo-) DNA polymerase (5u/µL) 1 µL
dd$H_2O$ 36.5 µL
$^{32}$P dCTP 5 µL
Incubate at 37 $^{o}$C for 2-10 minutes and add 2 µL stop solution
Pre-spin G-25 column for 1 minute at 735x g with lip loosen
Load 50 µL reaction product into the center of slope and spin at 735x g for 2 minutes
Count flow-through with program 27

3. Prepare 4% nature PAGE Gel:
Treat all parts in 0.1% NaOH overnight. Rinse all parts with tap water, clean with Windex followed by 70% ethanol, rinse with Millipore $H_2O$, and then dry. Wipe all parts with DNA/RNase removal reagent before use

50 mL 4% Gel:
30% Acrylamide:Bis (29:1) 6.67mL
10x TBE 1.25mL
10%APS 0.33mL
TEMED 40µL
dd$H_2O$ 41.7mL
Rinse wells and pre-Run gel in 0.5x TBE buffer at 100V, 1-2 hours

4. 2x mother mixture:
40% Glycerol and 2mM EDTA in 80mM Tris, pH 7.5 5 µL
KCL (1M) 1 µL
$MgCl_2$ (100mM) 1 µL
BSA (10mg/mL) 1 µL
DTT (20mM) 1 µL
Salmon sperm DNA (1mg/mL) 2 µL
Total Volume 11µL

5. Reaction:
2x Mother mixture 11µL
Nuclear extract 4µL
dd$H_2O$ 3µL
Block at RT for 5 minutes
*(For probe competition, add 50x cold probe and incubate at RT for another 5 minutes)*
$^{32}$P-probe 2µL
Incubate at RT for 15-30 minutes
Load 18 µL or all into gel
Run 2.5 hours at 150V at RT until the upper dye approaches 8 cm away from gel bottom

6. Dry gel:
Cover the gel with one piece of Whatman 3M paper and peer the gel off the plate
Stack the first Whatman paper on a second paper
Cover the other face of gel with Sara membrane and avoid bubbles
Transfer the gel within a radiation-protective container into gel dryer
Add ethanol and dry ice into the tank of gel dryer
Put the gel to the dryer with the paper facing to the bottom and cover the gel with dryer pad
Dry gel at 70 $^{o}$C for 1 hour

7. Exposure:
Pre-bleach the phosphor-storage plate
Replace the second Whatman paper with a third paper
Put the dried gel in the plate and cover for overnight
Scan the plate with Typhoon machine at 100-200 mp

**Bioinformatics and statistics.** RNA sequencing data analysis was performed with the pipelines in Basepair (New York, NY), GSEA 4.3.2 (11) and NIH David Function (12).

GEPIA tool (13) at http://gepia.cancer-pku.cn/detail.php was used to analyze the survival data in TCGA database. The Group cutoff was set as: High is >60% and low is <40%. NR2E3 RNA level was normalized by β-actin. *p* values were calculated by Logrank test.

NR2E3 gene expression profile in human tissues was obtained and sorted using GTEx online tool at https://gtexportal.org/home/gene/NR2E3#geneExpression (14).

NCBI ClinVar tool at https://www.ncbi.nlm.nih.gov/clinvar/?term=NR2E3%5Bgene%5D was used to overview SNVs of NR2E3.

The association study of NR2E3 mutations and cancer was done between TCGA and “All *of* Us” databases. “All *of* Us” database allows to stratify the population of > 182,000 cases with comprehensive information into sub-groups based on race, age and gender. Mutations that change NR2E3 protein sequence were collected in both databases. Chi-square test with two tails and 1 degree of freedom was used to calculate *p* values only of the comparisons with minimal case number of 5. Either Woolf logit test (> 99,000 cases) or Baptista-Pike (< 99,000 cases) test was used to calculate OR values with 95% confidence interval.

*p* values were calculated by Student’s t-test with two tails, except these individually described. The samples size was $n \geq 3$.

**Immunoblotting, Immunofluorescence, Co-Immunoprecipitation, RNA interference, Cycloheximide treatment, Reverse transcription followed by Real time-PCR, cell transfection, cell apoptosis and cell reporter assay** were previous published (1).

**Mutation primers:**

| Mutation | Primer sequence | |
|---|---|---|
| NR2E3-R76W | FWD | CAA GAG GAG CGT AtG GCG GAG GCT CAT C |
| | REV | GAT GAG CCT CCG CCa TAC GCT CCT CTT G |
| NR2E3-G88V | FWD | GCC AGG TGG GGG CAG tGA TGT GCC CCG |
| | REV | CGG GGC ACA TCa CTG CCC CCA CCT GGC |
| NR2E3-R97H | FWD | GAC AAG GCC CAC CaC AAC CAG TGC CAG |
| | REV | CTG GCA CTG GTT GtG GTG GGC CTT GTC |
| NR2E3-E121K | FWD | GCC GTG CAG AAC aAG CGC CAG CCG CG |
| | REV | CGC GGC TGG CGC TtG TTC TGC ACG GC |
| NR2E3-V302I | FWD | CAT GGA GAC GCG TaT CCT GCA GGA AAC |
| | REV | GTT TCC TGC AGG AtA CGC GTC TCC ATG |
| NR2E3-M407K | FWD | CTC CTT TGT GAT AaG TTC AAA AAC TGA ATT C |
| | REV | GAA TTC AGT TTT TGA ACt TAT CAC AAA GGA G |
| HA-NR2E3 short | FWD | GTT GTG TTA CCA TGT ATC CAT ATG ACG TCC CAG ACT ATG CCA TGG AGA CCA GAC CAA CAG CTC TG |
| | REV | GTT GTG AGC TCT CAC CTC ACG GGC TGG CTG GGG TG |
| HA-NR2E3-DBD | FWD | GTT GTG TTA CCA TGT ATC CAT ATG ACG TCC CAG ACT ATG CCA TGG AGA CCA GAC CAA CAG CTC TG |
| | REV | GTT GTG AGC TCT CAG GAC TCA GTG TTG GAC TCC ATG CTG |
| HA-NR2E3-LBD | FWD | GTT GTG TTA CCA TGT ATC CAT ATG ACG TCC CAG ACT ATG CCA TGC ATG AGA CCT CGG CTC GCC TAC TCT TCA TGG |
| | REV | GTT GTG AGC TCT CAG TTT TTG AAC ATA TCA CAA AGG AG |
| p53-L25-26A | FWD | CAG ACC TAT GGA AAg cAg cTC CTG AAA ACA AC |
| | REV | GTT GTT TTC AGG Agc Tgc TTT CCA TAG GTC TG |
| p53-R249S | FWD | CAT GAA CCG GAG tCC CAT CCT CAC |
| | REV | GTG AGG ATG GGa CTC CGG TTC ATG |
| p53-R273H | FWD | GAA CAG CTT TGA GGT GCa TGT TTG TGC CTG TCC TG |
| | REV | CAG GAC AGG CAC AAA Cat GCA CCT CAA AGC TGT TC |
| p53-R306A | FWD | CAG GGA GCA CTA AGg cAG CAC TGC CCA ACA ACA C |
| | REV | GTG TTG TTG GGC AGT GCT gcC TTA GTG CTC CCT G |

**ChIP-PCR and EMSA primers:**

| | | |
|---|---|---|
| p53BS1 on p21 (15) | FWD | GTG GCT CTG ATT GGC TTT CTG |
| | REV | CTG AAA ACA GGC AGC CCA AG |
| p53BS2 on p21 (15) | FWD | CCG AGG TCA GCT GCG TTA GAG |
| | REV | GCA GAG GAT GGA TTG TTC ATC |
| p53BS on MDM2 (16) | FWD | TGG GCA GGT TGA CTC AGC TTT |
| | REV | CCA GCT GGA GAC AAG TCA GGA |
| p53 EMSA probe | FWD | GAG TAC AGA ACA TGT CTA AGC ATG CTG GGG ACT |
| | REV | GTG AGT CCC CAG CAT GCT TAG ACA TGT TCT GTA |

**qRT-PCR primers:**

| Gene | Primer sequence | |
|---|---|---|
| p53BS1 on p21 | FWD | GCTGTGGCTCTGATTGGCTTT |
| | Probe | FAM-TGTCCCAAC |
| | REV | TTAGAGGTCTCCTGTCTCCTA |
| p53BS2 on p21 | FWD | ACAGCAGAGGAGAAAGAAGCC |
| | Probe | FAM-TGCGTTAGA |
| | REV | TCTCAGGCTCAGAGTCTG |
| p53BS on MDM2 | FWD | TGGGCAGGTTGACTCAGCTTT |
| | Probe | FAM-GTTCAGACACGTTCCGAAACTGCAGT |
| | REV | CCAGCTGGAGACAAGTCAGGA |
| UBE2L6 | FWD | CCACGGATGAGTCACAATCT |
| | REV | CCCAGGAACTGGCAATCTAA |
| EGR1 | FWD | CTCTACTGGAGTGGAAGGTCTA |
| | REV | GAACTTGGACATGGCTGTTTC |
| DHRS2 | FWD | GGTCTCTTCCATTGCAGCTTAT |
| | REV | CTCCAATGCCAGTGTTCTAGTG |
| IFI6 | FWD | GCTAGAGTGCAGTGGCTATT |
| | REV | GTAATCCTACTTGGGAGGTTGAG |
| IFI27 | FWD | CTGTCATTGCGAGGTTCTACT |
| | REV | ATTTGGGATAGTTGGCTCCTC |
| OAS1 | FWD | CAGTTGACTGGCGGCTATAA |
| | REV | TGTGAAGCAGGTGGAGAAC |
| OAS3 | FWD | GATGAGGGAGTGGGTCTATCT |
| | REV | TGGAGAGTCAGGCTGTCTAA |
| ABCB1 | FWD | CTTCATCGAGTCACTGCCTAAT |
| | REV | TAACAAGGGCACGAGCTATG |
| CA9 | FWD | GCTGTCTCGCTTGGAAGAA |
| | REV | TATTGGAAGTAGCGGCTGAAG |
| IGF1 | FWD | AACAAGCCCACAGGGTATG |
| | REV | ACATCTCCAGCCTCCTTAGA |
| CHAC1 | FWD | GGAGGCTTCTCTTTCTCAGTC |
| | REV | CACACCAACATGGTGCAATAA |
| UNC5B | FWD | CAAGGACAGTTACCACAACCT |
| | REV | CTGCCACTCCAAATGTGATAGA |
| ATF3 | FWD | CAGTTCCAAAGTCACAGGAAGA |
| | REV | CCTAGACACAACTCCTGACCTA |
| DDIT3 | FWD | AGGGAGAACCAGGAAACGGAAACA |
| | REV | TCCTGCTTGAGCCGTTCATTCTCT |
| TXNIP | FWD | CACTCTCAGCCATAGCACTTT |
| | REV | CATCTTCAGCCCACACTTTCT |
| TYMS | FWD | CAAATCTGAGGGAGCTGAGTAA |
| | REV | GAACAAAGCGTGGACGAATG |
| HDGF | FWD | TGGTCTCTCTATGCCTCTCTAC |
| | REV | ACGGTTCTCAGAGCTAAACTTC |
| CCNA2 | FWD | CTTCACCAGACCTACCTCAAAG |
| | REV | GGTGGGTTGAGGAGAGAAAC |

| | | |
|---|---|---|
| ACTB | FWD | CACTCTTCCAGCCTTCCTTC |
| | REV | GTACAGGTCTTTGCGGATGT |
| GAPDH | FWD | GCCTCAAGATCATCAGCAATGCCT |
| | REV | TGTGGTCATGAGTCCTTCCACGAT |
| B2M | FWD | TGTGTCTGGGTTTCATCCATCCGA |
| | REV | TCACACGGCAGGCATACTCATCTT |
| RPL38 | FWD | GCAGATACCTTTACACCCTGG |
| | REV | CTGGTTCATTTCAGTTCCTTCAC |

**Supplemental figure legends:**

**Figure S1: Gene-expression profiles of NR2E3.** (A) *NR2E3* is highly expressed in human urogenital systems (red font), thyroid, lung, *etc.*, besides in retinal photoreceptor cells. Data were extracted from GTEx Analysis Release V8 (dbGaP phs000424.v8.p2) at https://gtexportal.org/home/gene/NR2E3#geneExpression (14). (B) *Nr2e3* expression in mouse urogenital system and respiratory system was repeatedly detected. Data were extracted from Mouse Genome Informatics at https://www.informatics.jax.org/gxd/phenogrid/MGI:1346317 (17).

**Figure S2: NR2E3 selectively rescued the wild-type transactivities of p53 mutations which partially reserve this transactivity, but not the other mutations which completely lose it in p53-null HCT116 cells.** The indicated p53 plasmids was co-transfected with the p53 reporter and empty vector or NR2E3 plasmid into p53-null HCT116 cells. Two days later, the luminescence activity was quantified. The activity in the $p53^{WT}$+empty vector control was normalized to 1. *p* value was calculated by Student's t-test with two tails. *: $p<0.05$; **: $p<0.01$; ***: $p<0.001$. ***Notes:*** L25-26A is a mutation at the p53 TAD1 domain. R306A is a mutation at the p53 NLS domain. C135Y is a dominant-negative mutation. R249S is a hot-spot mutation. R273H is a gain-of-function mutation.

**Figure S3: Subfamily members of NR2E3 activate p53 transactivity.** NR2E3 and its subfamily members NR2E1, NR2F1 and NR2F2 that are all orphan nuclear receptors were co-transfected with the p53 reporter into HeLa cells (A) and HCT116 cells (B). The activity in control was normalized to 1. *p* value was calculated by Student's t-test with two tails. *: $p<0.05$; **: $p<0.01$; ***: $p<0.001$.

**Figure S4: Overview of Single Nucleotide Variants (SNV) of NR2E3.** The data were collected from ClinVar (https://www.ncbi.nlm.nih.gov/clinvar/?term=NR2E3%5Bgene%5D). (A) Classifications of NR2E3 SNVs. (B) Clinical involvements of NR2E3 SNVs.

**Figure S5: Summary of diseases-associated NR2E3 mutations whose molecular functions were reported in Enhanced S-Cone Syndrome.** *red font*: Pathogenic. R97H was detected in 2 out of 512 uterus endometrial cancer cases in a TCGA cohort.

**Figure S6: NR2E3 SNVs differentially regulate the p53 reporter.** NR2E3 mutations were co-transfected with the p53 reporter into in HCT116 cells (A) and H1299 cells (with $p53^{WT}$ plasmid) (B). Two days later, the luminescence activity was quantified. The activity in the mock control was normalized to 1. *p* value was calculated by Student's t-test with two tails. *: $p<0.05$; **: $p<0.01$; ***: $p<0.001$.

**Figure S7: Putative NR2E3 agonist-11a has a stronger inhibition of $p53^{+/+}$ HCT116 cells than $p53^{-/-}$ HCT116 cells.** The same number of $p53^{-/-}$ and $p53^{+/+}$ HCT116 cells were seeded in

6-well plates and the indicated concentrations of 11a were achieved in each well. Photos of these cells were taken daily from day-0 to day-5.

**Figure S8: 3-D spheroid culture of ATCC cervical cancer cell lines HeLa and SiHa.** (A-D) HeLa cells and (E-H) SiHa cells. (A) The combo eliminated spheroids from HeLa cells. The treatment started on day-6 when the spheroids were thriving. (B) The combo markedly decreased the number of HeLa spheroids on day-12. (C) The combo markedly suppressed the spheroid viability of MCW1 in CellTiter-Glo 3-D assay on day-12. (D) The combo stimulated the caspase 3/7 activity in MCW1 spheroids in Caspase-Glo 3/7 3D Assay on day-12. (E) The combo eliminated spheroids from SiHa cells. (F) The combo markedly decreased the number of SiHa spheroids on day-15. (G) The combo markedly suppressed the spheroid viability of SiHa on day-15. (H) The combo stimulated the caspase 3/7 activity in SiHa spheroids on day-15. *p* value was calculated by Student's t-test with two tails. *: $p<0.05$; **: $p<0.01$; ***: $p<0.001$.

**Figure S9: 11a specifically enhances NR2E3-stimulated p53 transactivation in HeLa cells.** (A) 11a activated the p53 reporter much more in NR2E3 group than other groups. NR2E3 or its subfamily members were co-transfected with the p53 reporter into HeLa cells in 96-well plates. One day after transfection, 11a was added to achieve the indicated concentrations in each well. One day after 11a treatment, the luminescence was measured. The activity in the DMSO control in mock control was normalized to 1. (B) NR2E3 shRNAs disrupted the 11a's stimulation of the p53 reporter. The indicated shRNAs were co-transfected with the p53 reporter into HeLa cells. One day after transfection, 11a was added to achieve the indicated concentrations in each well. One day after 11a treatment, the luminescence was measured. The activity in the DMSO and scramble shRNA control was normalized to 1. *p* value was calculated by Student's t-test with two tails. *: $p<0.05$; **: $p<0.01$; ***: $p<0.001$.

**Figure S10: 11a induces significant transcriptome changes in HeLa cells.** RNA sequencing was conducted in HeLa cells treated with 11a using the conditions of Figure 5C. (A) Volcano plot of all transcripts of 11a and DMSO control. (B) Heat map of differentially expressed genes. (C) Changes of these selected genes were verified using qRT-PCR. (D) Overview of GSEA results labeled with gene sets. (E) p53, apoptosis, and TGFβ pathways were enriched by 11a. (F) Oncogenic E2F and PI3K pathways and (G) Metabolic glycolysis and oxidative phosphorylation pathways were enriched in DMSO controls. (H) Oncogenic TNFα and KRAS pathways were enriched in 11a treated samples. *p* value was calculated by Student's t-test with two tails. *: $p<0.05$; **: $p<0.01$; ***: $p<0.001$.

**Figure S11: Summary of Hallmark GSEA data in the comparisons of 11a vs DMSO, Romidepsin vs DMSO, Combo vs DMSO, Combo vs 11a, and Combo vs Romidepsin.** Not enriched: FDR > 0.25.
